# Molecular mechanism of partially open Kv7.2 gate stabilization by gain-of-function variants

**DOI:** 10.64898/2026.07.15.738424

**Authors:** Agnese Roscioni, Giulio Alberini, Francesco Miceli, Fabio Benfenati, Maurizio Taglialatela, Luca Maragliano

## Abstract

Gain-of-function (GoF) variants in the Kv7.2 channel are associated with a clinically relevant subset of neurodevelopmental disorders. While most GoF substitutions identified so far affect the voltage-sensing domain, we recently described three mutations in the intracellular-facing activation gate (AG), G313S, A317T, and L318V. Electrophysiological recordings showed that these variants increase macroscopic current density and enhance channel open probability. Consistently, molecular dynamics (MD) simulations revealed that they hinder complete channel closure by destabilizing the closed AG and increasing hydration of the central cavity (CC). Whether this partially open conformation can support K^+^ permeation, however, remained an open question. Here, we combined long-timescale atomistic simulations and simulations with applied electric fields to evaluate the stability of these mutant-associated AG states over longer timescales and their functional relevance. In new trajectories, all three substitutions consistently shifted the closed intracellular gate toward a widened, water-accessible conformation, accompanied by increased CC hydration. We then assessed the functional significance of this partially open state by simulating the A317T channel under applied electric fields. The conformation supported K^+^ translocation, whereas the closed WT pore remained impermeable under all tested voltages. When simulations were started from open channel conformations, both WT and A317T conducted K^+^ ions, indicating that the main effect of the substitution is to destabilize closure of the intracellular gate rather than to alter the fully conductive open state. Together, these data show that AG GoF variants can generate an intermediate gate conformation that permits ion permeation, providing a mechanistic link between mutant-induced pore remodeling and Kv7.2 dysfunction in KCNQ2-related disease.

## Introduction

The neuronal members of the Kv7 voltage-gated potassium channel family, Kv7.2 to Kv7.5, contribute to the M-current, a slowly activating, non-inactivating potassium current that plays a key role in tuning neuronal excitability (Hoshi 2020; Jepps et al. 2021; Dirkx et al. 2020; Alberini et al. 2021; Zuberi et al. 2022). Pathogenic variants in Kv7.2, encoded by the KCNQ2 gene, are associated with a spectrum of neurodevelopmental disorders, including encephalopathies with or without epilepsy, making this protein a valid pharmacological target for the design of new drugs (Weckhuysen et al. 2012; Goto et al. 2019; Malerba et al. 2020; Nappi et al. 2022, 2024). Structurally, Kv7.2 can form homotetrameric channels, with the four subunits symmetrically arranged around a central ion-conducting pore. Each subunit comprises six transmembrane (TM) α-helices (S_1_ to S_6_): the first four form the voltage sensor domain (VSD), while S_5_-S_6_, together with the pore loop, define the pore domain (PD). The latter includes the selectivity filter (SF), facing the extracellular side, and the activation gate (AG, also named inner gate), facing the intracellular side, separated by the central cavity (CC).

While most Kv7.2 variants, when expressed *in vitro*, reduce channel function by a variety f mechanisms, and are therefore referred to as “loss-of-function” (LoF) variants, few of them enhance channel function, producing gain-of-function (GoF) *in vitro* phenotypes (Nappi et al. 2020); these latter variants are mainly localized in the VSD (Miceli et al. 2015; Miceli et al. 2022, Millichap et al. 2017). Instead, three GoF substitutions, G313S, A317T, and L318V, were recently identified in the TM S_6_ helix, which is part of the AG (Nappi et al. 2022, 2024). Electrophysiological recordings showed that the mutations increase the macroscopic current density, enhance the channel open probability, and cause an instantaneous, voltage-independent current component compared with wild-type (WT) Kv7.2. Consistently, molecular dynamics (MD) simulations suggested that, when introduced into a channel with a closed AG, these substitutions promote a broadening of the gate and an increased hydration of the CC, whereas simulations initiated from an open AG revealed no major differences between WT and mutant channels (Nappi et al. 2024). These results support a model in which the variants favor an incomplete closure of the AG, which may represent a mechanistic signature of GoF-associated Kv7.2 encephalopathies. However, whether the MD-derived conformation corresponds to a pore architecture compatible with ion permeation remains unresolved. Addressing this question requires testing both the structural robustness of the variant-driven rearrangement and the ability of the resulting pore configuration to sustain K^+^ permeation. Here we performed new MD simulations of the three variants over longer timescales (up to 10 μs for one system) and tested whether the structural rearrangement was reproduced with two different force fields, CHARMM (Huang et al. 2017) and AMBER (Maier et al. 2015). In addition, we probed ion permeation under applied electric fields (Crozier et al. 2001; Aksimentiev et al. 2004; Gumbart et al. 2012) in A317T Kv7.2, the variant associated with the most pronounced electrophysiological phenotype (Nappi et al. 2024). CHARMM-based simulations of all variants, initiated from a closed AG, confirmed the stabilization of a partially open gate, associated with enhanced hydration of the CC. By contrast, AMBER trajectories revealed less pronounced differences between mutants and WT, with the pore remaining closer to the native closed-state geometry. We then used CHARMM to test whether incomplete AG closure can support ion conduction. Applied-field simulations of A317T showed that the partially open conformation permits K^+^ permeation, while the WT closed structure remains non-conductive. Conversely, both WT and A317T open structures displayed significant K^+^ translocation, confirming that the mutation primarily affects the closed-state gate architecture. Overall, these results provide a structural and functional characterization of the pore rearrangements induced by disease-associated KCNQ2 GoF variants, offering new mechanistic insight into their contribution to neurodevelopmental disorders.

## Materials and Methods

### Choice and characterization of channel structures

The simulations described in this work were performed on the human Kv7.2 channel, considering the WT and mutant variants in different conformational states. Following our previous work (Nappi et al. 2024), we used five cross-distances (CDs) to distinguish among channel conformations. The first distance (d1) defines the conductance state of the SF, while the other four (d2 to d5) describe the degree of AG opening (Suppl. Fig. S1). Each CD is defined between two identical atoms of diagonally opposed subunits, and is therefore computed twice (between chains A and C and between chains B and D):

- d1, between the Cα atoms of two G279,
- d2, between the Cα atoms of two G313,
- d3, between the Cα atoms of two A317,
- d4, between the Oγ atoms of two S314,
- d5, between the Cδ atoms of two L318.

#### Closed AG structure

To represent the closed-gate state, we used the apo cryo-EM model from Li et al. 2021(PDB ID: 7CR0). This conformation is characterized by a conductive SF (d1= 8.03Å) and an occluded AG (d2= 11.67 Å, d3= 13.69 Å), with two distinctive constrictions at the level of S314 and L318 (d4= 4.00 Å, d5 = 7.00 Å). MD simulations were performed using only the pore domain (PD, residues G215-E330), omitting the four voltage-sensor domains (VSDs). USFC Chimera (Pettersen et al. 2004) was employed to insert the mutations at the corresponding sites (i.e., G313S, A317T, L318V), based on the Dunbrack rotamers library (Dunbrack Jr and Karplus 1993).

#### Open AG structure

For the open-gate conformation, we used two different structures, each characterized by distinct CD values:

- The cryo-EM structure of the cannabidiol-activated channel (PDB ID: 8J01, (Ma et al. 2023)), with d1= 8.05 Å, d2=13.77 Å, and d3=18.23.
- A homology model (HM) based on the Kv7.1 open cryo-EM structure (PDB ID: 6V01) (Sun and MacKinnon 2020), previously designed and employed by us in Ref. (Nappi et al. 2024), and characterized by d1=7.97 Å, d2=13.71 Å, and d3=19.43 Å.

### Equilibrium molecular dynamics simulations

The closed conformations of Kv7.2 WT and G313S, A317T, L318V variants were simulated using the GPU-enabled GROMACS 2023 package (Bekker et al. 1993, Berendsen et al. 1995, Abraham et al. 2023) and two alternative force-field parameterizations, CHARMM36m and AMBER. A summary of the simulations conducted is provided in Table 1.

**Table 1:**
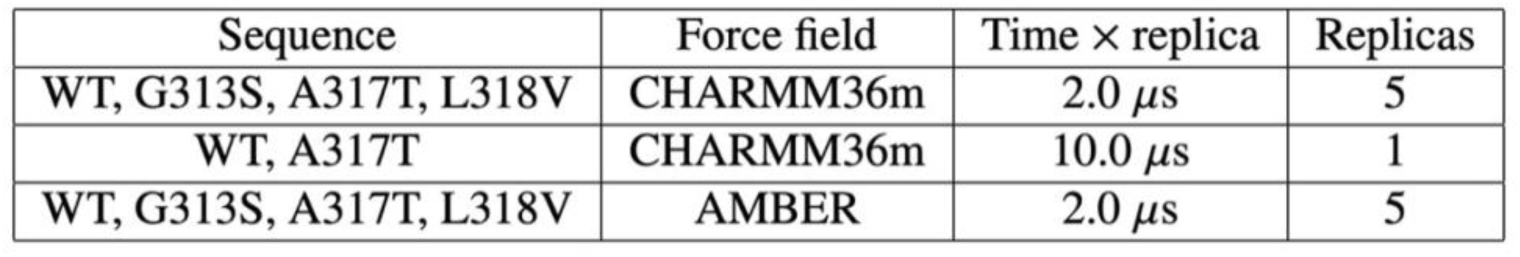
Summary of equilibrium MD simulations of WT and mutant Kv7.2 channel (starting structure: PDB ID 7CR0, Li et al. 2021, closed AG). Cumulative simulation time: 100 µs.

#### Systems preparation

The CHARMM-GUI (C-GUI) Membrane Builder server (Jo et al. 2007) was used to prepare all necessary simulation input files. Each of the four Kv7.2 pore structures (WT and G313S, A317T, and L318V variants) was oriented within the membrane using the OPM-PPM server (Lomize et al. 2012) and then inserted into a homogenous 1-palmitoyl-2-oleoyl-sn-glycero-3-phosphocholine (POPC) membrane. The total charge of each system was neutralized with a physiological 150 mM KCl solution, yielding approximately 155,000 atoms.

#### CHARMM settings

The CHARMM36m and CHARMM36 parameter sets were used for the protein and lipids (Huang et al. 2017; Best et al. 2012; Huang and MacKerell 2013), respectively, together with the CHARMM36-modified TIP3P water model (Jorgensen et al. 1983) and CHARMM-compatible ion parameters with NBFIX corrections (Noskov and Roux 2008; Venable et al. 2013; Luo and Roux 2010). Periodic boundary conditions (PBCs) were applied, and long-range electrostatic interactions were computed using the Particle Mesh Ewald (PME) method (Darden et al. 1993). The cutoff for non-bonded interactions was set at the CHARMM standard of 12 Å, with a smooth force-switching function between 10 and 12 Å. Covalent bonds involving hydrogen atoms were constrained with LINCS (Hess et al. 1997), allowing an integration timestep of 2 fs. Systems equilibration adhered to the standard C-GUI membrane builder protocol (Jo et al. 2007), involving multiple steps with progressive relaxation of positional restraints. This was followed by an additional equilibration run of 60 ns under the last-stage conditions (weak restraints on protein backbone heavy atoms only). After equilibration, all simulations were performed in the NPT ensemble at 310 K using the suggested inputs for production, including the Nose-Hoover thermostat (coupling constant of 1.0 ps) (Nosé 1984; Hoover 1985) and at 1 atm with the Parrinello-Rahman barostat (coupling constant of 5.0 ps) (Parrinello and Rahman 1981) with semi-isotropic pressure coupling. For each of the four systems, five independent replicas of 2 *μ* s each were simulated. Additionally, the WT and A317T systems were simulated up to 10 *μ*s to explore longer-timescale events.

#### AMBER settings

These simulations were performed using the AMBER14SB (Maier et al. 2015) and LIPID21 (Dickson et al. 2022) parameter sets. The systems were solvated with TIP3P water (Jorgensen et al. 1983) and Joung and Chatman ion parameters (Joung and Cheatham III 2008). The cutoff for non-bonded interactions was set at 9 Å. All other parameters and protocols were as described above for the CHARMM settings. For each of the four systems, five independent replicas of 2 *μ*s were simulated.

### Molecular dynamics simulations with a constant external electric field

We simulated Kv7.2 WT and A317T in closed/partially open and open-gate conformations under a constant, uniform external electric field applied normal to the membrane. The NAMD code (Phillips et al. 2005) was used for this purpose. Since in our previous NAMD simulations we used the CHARMM36 force-field (Nappi et al. 2024), we regenerated the systems using C-GUI and repeated the full setup and equilibration from scratch to employ the updated CHARMM36m parameters. Again, only the protein PD was considered (residues G215-E330).

#### Systems setup and equilibration

Systems were prepared using C-GUI as described above, except that now a 500 mM KCl solution was employed. Simulations were performed using the CHARMM36m/CHARMM36 force field for protein and lipids, respectively (Huang et al. 2017; Huang and MacKerell 2013; Klauda et al. 2010), together with the CHARMM-modified TIP3P water model (Jorgensen et al. 1983) and CHARMM-compatible ion parameters with NBFIX corrections (Noskov and Roux 2008; Venable et al. 2013; Luo and Roux 2010). Tetragonal PBCs were applied to the simulation box to remove surface effects, and the PME method was used to calculate long-range electrostatic interactions (Darden et al. 1993). Short-range electrostatic and van der Waals interactions were calculated with a 12-Å cutoff, with a smooth decaying function starting to take effect at 10 Å. The CHARMM force-based switching function was employed for van der Waals interactions via the vdwForceSwitching command (Steinbach and Brooks 1994). The SHAKE algorithm (Ryckaert et al. 1977) constrained covalent bonds involving hydrogen atoms (except for water molecules, where SETTLE (Miyamoto and Kollman 1992) was used), allowing an integration time step of 2 fs. To ensure maximum accuracy, electrostatic and van der Waals interactions were computed at each simulation step. We used the Nosé-Hoover Langevin piston method (Feller et al. 1995; Martyna et al. 1994) and a Langevin thermostat to reproduce the NPT ensemble, maintaining pressure at 1 atm and temperature at 310 K. According to the C-GUI input files (Jo et al. 2007), the oscillation period of the piston was set at 50 fs, with a damping time scale of 25 fs. The Langevin thermostat was set with a damping coefficient of 1 ps^-1^. For the NVT equilibration, the barostat was switched off, and the Langevin thermostat was used to maintain the temperature constant at 310 K. In some of the equilibration stages described below, we used the Hydrogen Mass Repartitioning (HMR) approach, with a time step of Δ*t* = 4 fs (Feenstra et al. 1999; Hopkins et al. 2015; C. Balusek et al. 2019). For HMR simulations, we fixed the piston period at 300 fs and and the langevin piston decay at 150 fs (Curtis Balusek et al. 2019).

Before applying the electric field, systems were equilibrated via a two-stage protocol (NPT followed by NVT), with different procedures for the initially closed- and open-gate conformations, summarized here and reported in detail in the supplementary file (Supplementary Section 1) and Suppl. Figs. S2 and S3. **Initially closed structures:** the NPT stage comprised an extended C-GUI equilibration, a trajectory of 1 *μ* s with HMR, and subsequently, 250 ns with standard masses. In this phase, dihedral restraints were applied to the SF (residues 276-283) to prevent collapse. Then, for both systems, a conformation was extracted from the last 200 ns using the GROMOS clustering algorithm (Daura et al. 1999), after verifying that pore opening and hydration were consistent with our previous simulations (Nappi et al. 2024). These structures were further equilibrated in NVT for 50 ns, with positional restraints that were gradually removed and boundary restraints on CDs d2 and d3. Finally, the external electric field was applied in successive 10 ns runs by increasing the transmembrane potential in 200 mV increments from 100 mV, until each target value (see below) was reached. **Open structures:** the NPT stage included an extended C-GUI equilibration, followed by additional 500 ns with lower-bound distance and SF restraints to allow proper hydration of the channel. In the subsequent NVT phase, we followed the same procedure just described for the initially closed AG structures.

#### Constant electric field protocol

Starting conformations for closed/partially open and open WT and A317T were obtained from the equilibration stages described above. Typical box dimensions were ∼[120 Å × 120 Å × 114 Å.

Simulations were run in NVT conditions, with temperature maintained by a Langevin thermostat applied only to lipid atoms (damping 1 ps^−1^), while all other atoms were left unthermostatted (zero damping), so as to dissipate any potential field-induced heating without directly biasing ion dynamics (Sotomayor et al. 2007).

A constant uniform electric field *E_z_* was applied along the membrane normal *z*, corresponding to an imposed transmembrane voltage *V* = *E_z_L_z_* across the periodic simulation cell, where *L_z_* is the box length along the *z* direction (Crozier et al. 2001; Aksimentiev et al. 2004; Gumbart et al. 2012). We chose the constant electric field approach over the double-bilayer method (Kutzner et al. 2011; Machtens et al. 2015) to reduce system size and computational cost.

The following restraints were applied to prevent structural distortion of the proteins:

- Positional restraints on the C*α* atoms of transmembrane (TM) helices (i.e., residues 238-247 and 294-303 of each helix), in all systems.
- Lower- ad upper-bound restraints on the d2 and d3 distances, based on values from previous standard MD simulations (Nappi et al. 2024). WT system: closed AG, d2 between 11.80 Å and 12.20 Å, and d3 between 11.80 Å and 12.30 Å; open AG, d2 between 15.04 Å and 15.76 Å, and d3 between 18.05 Å and 19.29 Å. A317T: partially open AG, d2 between 11.50 Å and 13.50 Å and d3 between 13.00 Å and 16.00 Å; open AG, d2 between 14.92 Å and 15.92 Å and d3 between 18.23 Å and 19.65 Å.
- Lower-bound restraints on the inter-subunit distances between C *α* atoms of residues R325 to Q330, to keep them larger than 35 Å and prevent the truncated, flexible C-terminal segments from occluding the AG (all systems).
- To avoid the collapse of D282 over the SF observed in simulations initiated from the open-gate Cryo-EM structure, only for this system we applied a lower-bound restraint to the inter-subunit distances between C*α* atoms of G281 residues to keep them larger than 8.10 Å.

Different voltages were applied to each system. To obtain an appreciable number of ion permeation events within a computationally accessible timescale (500 ns), we employed supraphysiological voltages, ranging from -1.3V to 1.7V (Machtens et al. 2015). Throughout all simulations, the proteins remained structurally stable also in unrestrained regions, and no lipid bilayer poration was observed. Seven replicas were run for each voltage value. For the open systems, four replicas used the cryo-EM structure, and three replicas our homology model. In Table 2, we report a summary of the simulated systems.

**Table 2:** Summary of constant electric field simulations of the Kv7.2 channel. Cumulative simulation time: 84 µs.

| Sequence | AG | Starting Structure | Voltage values (V) | Force field | Time<br>× replica | Replicas<br>per voltage |
| --- | --- | --- | --- | --- | --- | --- |
| WT | Closed | PDB 7CR0 | -1.3; 0.9; 1.1;<br>1.4; 1.5; 1.7 | CHARMM36m | 500 ns | 7 |
| A317T | Partially open | PDB 7CR0 | -1.3; 0.9; 1.1;<br>1.4; 1.5; 1.7 | CHARMM36m | 500 ns | 7 |
| WT | Open | PDB 8J01 / HM | -1.3; 0.9; 1.1;<br>1.4; 1.5; 1.7 | CHARMM36m | 500 ns | 7 |
| A317T | Open | PDB 8J01 / HM | -1.3; 0.9; 1.1;<br>1.4; 1.5; 1.7 | CHARMM36m | 500 ns | 7 |

### Analysis of ion passage and configurations within the filter

#### Ion counting

Ion permeation events were counted with an in-house TCL script. Two reference surfaces (disks) were defined, one at the entrance of the AG (disk1), one at the extracellular end of the SF (disk2). A permeation event was recorded when an ion crossed disk1 and subsequently disk2 (the reverse order for the negative voltage).

#### Persistence of ionic configurations

Configurations of ions within the filter were characterized by evaluating, for each frame of the seven replicas, the occupancy of the four coordination sites. Each site was labeled as water (W), potassium (K), or void (0), forming a four-character string per frame. Configurations were then associated with the minimal d1 value in that frame (d1-min). The persistence of each state in all replicas is reported as a percentage.

### Analysis and visualization of the molecular structures

The COLVARS (Collective Variable-based Calculation) module (Fiorin et al. 2013) was employed to calculate distances and other observables, and to impose restraints. Analyses were also performed with VMD, Visual Molecular Dynamics (Humphrey et al. 1996), and UCSF Chimera and ChimeraX (Pettersen et al. 2004, 2021; Goddard et al. 2018; Meng et al. 2023). For simulations performed with the GROMACS code, root-mean-square distances (RMSDs), root-mean-square fluctuations (RMSFs), and distance calculations were performed using functions integrated into GROMACS (respectively, rms, rmsf, mindist). PROLIF (Bouysset and Fiorucci 2021) and MDAnalysis (Michaud-Agrawal et al. 2011) were used for interaction calculations. Calculations of the channel profile along the main axis were performed using HOLE (Smart et al., 1996, 1997). Hydration analysis along the pore axis was also performed as in our previous study (Nappi et al. 2024). Pictures were created with ChimeraX.

## Results and Discussion

### Equilibrium molecular dynamics simulations

To assess variant-induced effects on the closed-AG Kv7.2 structure over extended timescales and with different force fields, we performed five independent 2 *μ* s MD simulations of WT and the G313S, A317T, and L318V mutants, using both CHARMM and AMBER parameter sets and the TIP3P water model. In addition, two 10*μ*s-long trajectories were generated for WT and A317T using CHARMM.

#### CHARMM force-field

The time evolution of the backbone RMSDs for WT (Suppl. Fig. S4A) and each variant (Suppl. Fig. S4B-D) showed stable plateaus around 2-2.5 Å for both the tetramers and individual subunits, supporting the structural stability of the PD even when deprived of the VSDs, and indicating that each variant preserves the overall WT fold. We then looked for structural differences in more detail, starting with the pore radius profile. Fig. 1 shows, for each system, the pore radius averaged over all MD replicas. Compared with WT, each mutant exhibited a widening of the closed-AG region and, for G313S and A317T, also of the central cavity. Notably, all three mutants showed a loosening of the two constrictions distinctive of the closed conformation, at residues 314 and 318. Consistently, the distances d4 and d5 were larger in the mutants than in the WT throughout the simulations (Suppl. Fig. S5). This broadening led to increased pore hydration, as indicated by the average number of water molecules along the channel axis (Fig. 1C, F, I).

**Figure 1:**
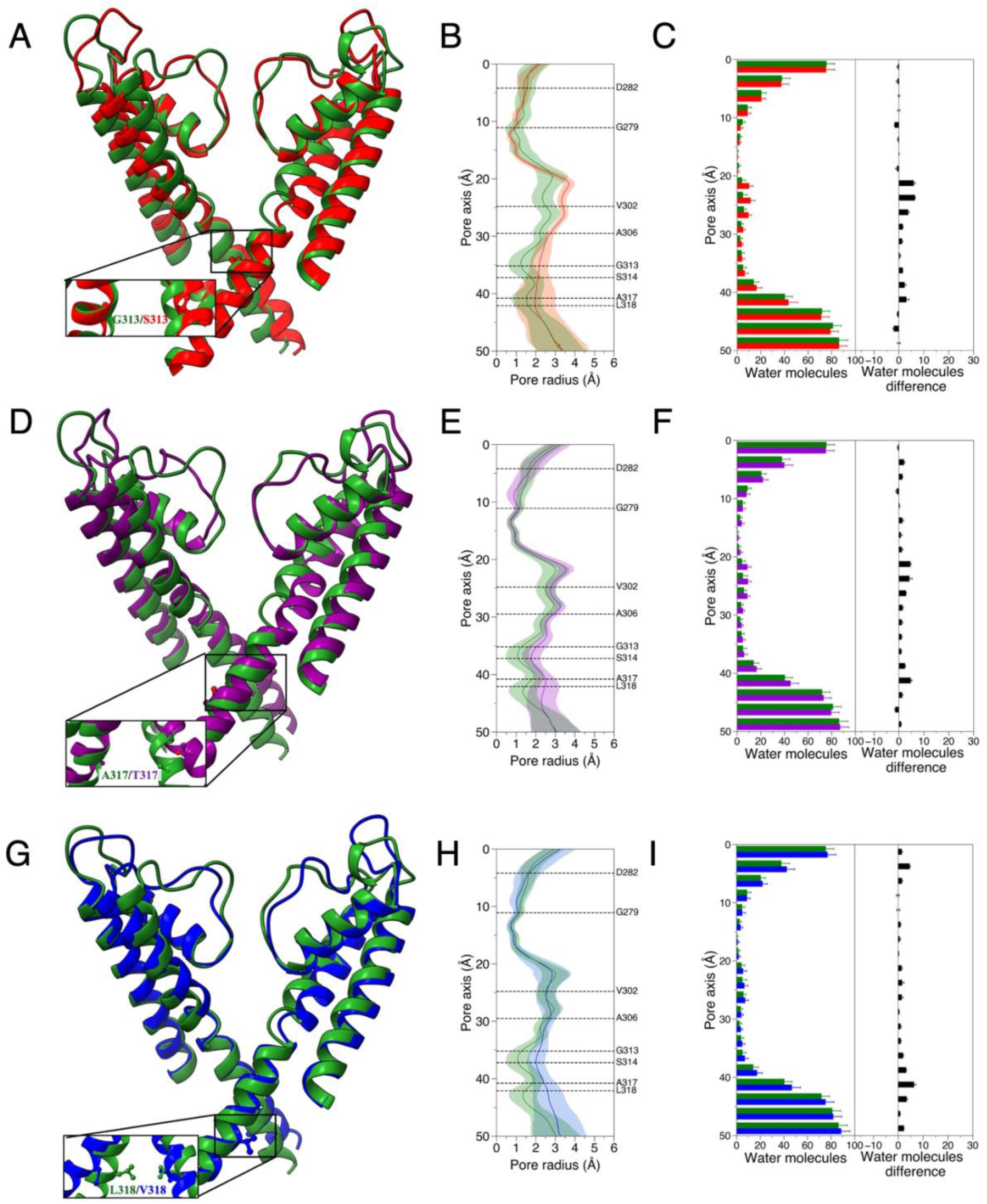
CHARMM MD simulations of WT and mutated channels, starting from the closed AG. (A, D, G) Superposition of WT (green) and variant structures at the end of 2 μs trajectories: G313S (red), A317T (purple), and L318V (blue). (B, E, H) Channel pore radius profiles averaged over all simulated replicas. Shaded regions represent the associated standard deviations. (C, F, I) Distribution of water molecules within the channel pores and differences between mutant and WT (panels on the right). Averages and standard deviations are calculated using all replicas.

Interestingly, in all systems, the conductive state of the SF was preserved: Suppl. Fig. S6 shows the RMSFs of the SF residues and the evolution of the d1 CD, confirming the stability of the filter.

To explore mutation effects on even longer timescales, we performed two 10 *μ*s simulations starting from closed conformations of Kv7.2 WT and A317T, focusing on this variant because it yielded the most pronounced experimental effect (Nappi et al. 2024). Results were consistent with those from shorter simulations. The WT channel showed again high stability, with the protein backbone RMSD stable around ∼ 2.5 Å (Fig. S7A), d1 close to the conductive-filter threshold, and d4 and d5 always near the closed-AG values, showing only minor fluctuations (Suppl. Fig. S7C). Conversely, A317T data revealed a conformation change: the backbone RMSD stabilized between 3 Å and 3.5 Å (Suppl. Fig. S7B) and, while d1 remained consistent with a conductive SF, d4 and d5 explored much larger values (Suppl. Fig. S7D). Calculation of the pore profile and hydration indicated widening at both the AG and the CC (Suppl. Fig. S7E,F), and a larger number of water molecules within the mutant channel (Suppl. Fig. S7G). Overall, the results of this section confirm what was observed in our previous work using shorter timescales (Nappi et al. 2024), i.e., that the three mutants destabilize the channel closed configuration at the level of the AG, inducing pore widening and increased hydration, while leaving the filter in the conductive conformation.

As in our earlier work, we then searched for interactions that could be responsible for the observed structural modifications. In the G313S system (Suppl. Fig. S8 A,B), we again observed a highly persistent intra-chain H-Bond between S313 and A309, which induced a tilt and rotation of each S_6_ helix. As a consequence, the S314 side chains, one of the characteristic closed-AG constrictions, are displaced away from the pore lumen. The orientation of S313 is further stabilized by a persistent H-bond with A317. In the A317T system (Suppl. Fig. S8 C-E), we observed the previously identified H-bond between the T317 and S314 residues of adjacent chains, again displacing the side chains of the closed-AG constriction. The substituted residue is also stabilized by an intra-chain interaction with G313, and by an H-bond between the backbone carbonyl oxygen and the Q321 amide group. For L318V (Suppl. Fig. S8F,G), we did not find specific interactions associated with pore widening via helix rearrangement or side chain displacement. Instead, the reduced steric bulk and hydrophobic surface area of the substitution led to a loosening of the constriction at position 318, facilitating water entry and promoting upward propagation of the dilation.

#### AMBER force-field

To evaluate the sensitivity of the variant-induced effects to the choice of force field, we repeated the 2 *μ*s simulations of WT and the three variants using AMBER (version ff14SB), again running five replicas per system. The backbone atoms RMSD of the WT channel was stable for both the whole protein and single subunits (Suppl. Fig. S9A). The variants’ RMSDs showed larger fluctuations for some chains (Suppl. Fig. S9 B-D), and, for A317T, a modest drift. These differences mapped onto distinct pore features in the three mutants (Suppl. Fig. S10): all variants showed more localized gate widening, and L318V exhibited a constriction of the central cavity; CC hydration increased in G313S, remained unchanged in A317T, and decreased in L318V. As for the gate CDs d4 and d5 (Suppl. Fig. S11), both are moderately larger and more variable in the mutants than in WT. Finally, the conductive conformation of the SF is also preserved in these simulations, as revealed by residue RMSFs and by the time evolution of the d1 CD (Suppl. Fig. S12). These results indicate that, at least for the explored timescales, mutation-induced structural deviations are generally less pronounced in the AMBER simulations, with the pore remaining closer to the WT geometry than in the corresponding CHARMM simulations. When looking at the interaction patterns (Suppl. Fig. S13), we found that, while the key, variant-induced H-bonds are present, additional interactions form that might compensate and hinder the conformational change.

### Constant electric field simulations

Next, we set out to assess whether the mutation-driven widening of the closed AG renders the pore conductive. To this end, we performed MD simulations under constant external electric fields, applied along the membrane normal. We focused on the A317T variant because it showed the most pronounced experimental effect, and simulated the partially open mutant and the closed WT conformations, as well as the open WT and variant structures. The NAMD code and the CHARMM36m force field were employed.

#### Partially open A317T and closed WT

Starting conformations for the constant field trajectories were obtained after preliminary equilibration in NTP and NVT, as described in Materials and Methods and in supporting material (see Suppl. Fig. S2 for a summary), during which the backbone RMSDs were stable around 2-2.5 Å for both systems (Suppl. Fig. S14). Once again, starting from the closed AG structure, the mutant displayed a widened pore and a hydrated CC (Suppl. Fig. S15). To monitor SF stability along the applied voltage trajectories, we calculated four distances between carbonyl oxygens of equivalent residues across subunits, named d_s1 to d_s4, starting from the extracellular side (Suppl. Fig. S16). At all voltages, the four coordination distances did not differ substantially from the cryo-EM reference, except for the outermost d_s1, likely because of its location in the relatively more flexible region of the filter.

At each transmembrane voltage (-1.3V to 1.7V), the partially open A317T channel displayed multiple K^+^ translocation events across the pore, traversing the AG and the SF, consistent with voltage-driven conduction. In Fig. 2, we show, for representative replica trajectories at three selected voltages, the time evolution of ions and water *z*-coordinates within the filter, together with the boundaries of the ion binding sites defined by the centers of mass of opposing carbonyl groups (Khalili-Araghi et al. 2006; Kopec et al. 2019; Ocello et al. 2020). The d1 CD values are reported below each panel, to monitor again filter stability during permeation events. Similar plots for the other replicas at the same voltages are provided in Suppl. Figs. S17 to S19. By contrast, the closed WT channel showed no permeation events at any tested voltage (Suppl. Figs. S20–S22), with the SF showing only short-lived narrowing, as revealed by the d1 traces.

**Figure 2:**
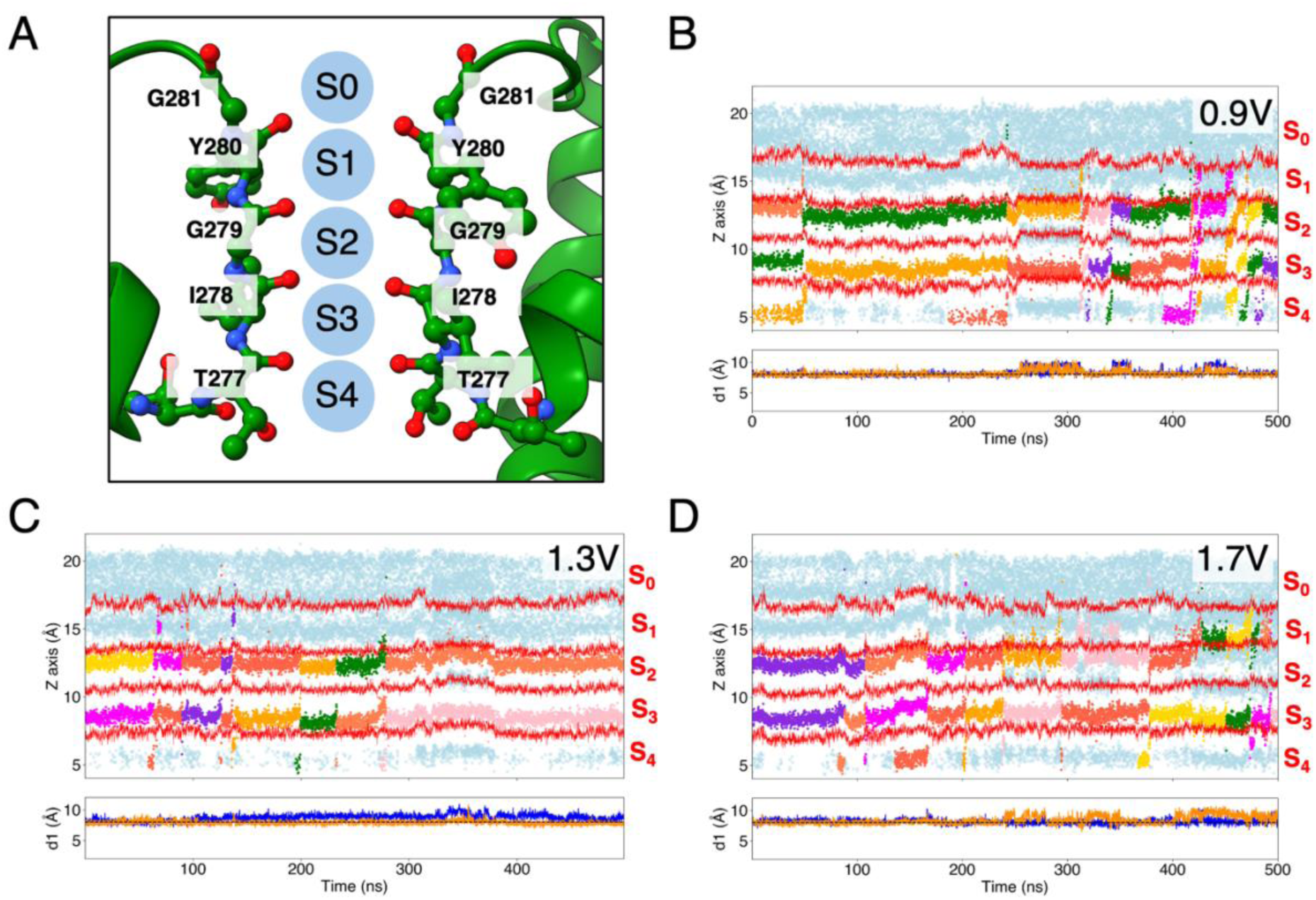
Ion translocation through the A317T SF in applied electric field simulations. (**A**) 3D representation of the SF and the coordination sites. (**B**-**D**) Top panels: time evolution of the z-coordinates of K^+^ ions (different colors), water (light blue) and SF residues (red) at three different transmembrane potentials; bottom panels: time evolution of the d1 distance (calculated between protein chains A-C (blue) and B-D (orange)) during the same trajectories, the dashed black line shows the reference d1 value (8.03 Å) from the PDB ID: 7CR0 cryo-EM structure.

In Fig. 3, we report the time series of cumulative K^+^ permeation events through the closed WT and partially open A317T, calculated at all transmembrane voltages considered. While the native channel shows no translocation at any voltage (black trace), the mutant supports sustained K^+^ flux, with the cumulative permeation count increasing monotonically over time. These results indicate that the mutation-driven broadening of the closed AG is compatible with ion permeation, consistent with electrophysiology measurements showing non-negligible mutant currents under conditions where the WT is silent (Nappi et al. 2024).

**Figure 3:**
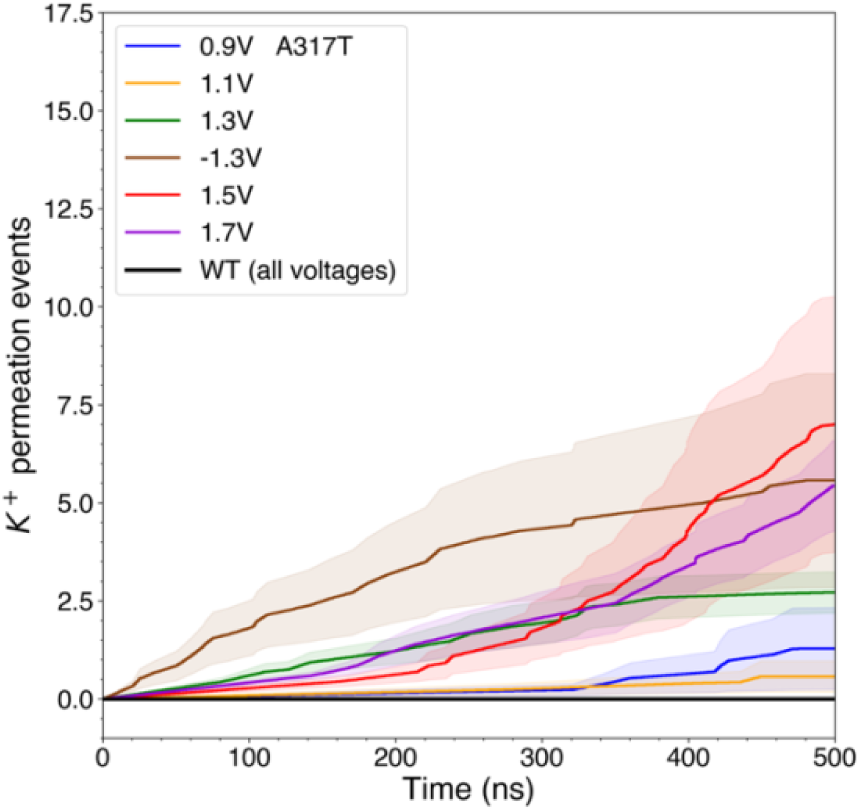
Time evolution of K^+^ permeation events through the partially-open variant channel structures, averaged across replicas at each voltage.

Having established K^+^ conduction in the mutant channel, we next characterised the permeation mechanism by quantifying the occupancies of the SF site and the resulting ion-water configurations sampled along the applied field trajectories. To this aim, we indicate the occupancy state as a string of four characters, each corresponding to one of the S1 to S4 binding sites, chosen between K, W, or 0 depending on whether each site is occupied by a potassium ion, a water molecule, or is empty, respectively. Statistics of the occurrence of SF states were accumulated over all replicas and voltages and correlated with the corresponding filter width by considering the smallest of the two d1 values (indicated as d1-min). Results are reported in Fig. 4 as violin plots, showing the occurrence percentage of each configuration and the corresponding distribution of d1-min, and summarized in Table 3. In the partially open A317T mutant at the negative voltage, the two most frequent configurations are 00WK and W0WK, both associated with a restricted filter, with W0WK corresponding to the more constricted state. At all positive voltages, the most frequent conformation is WKK0, and the second is WKKK, except for 1.5 V, where it is WWK0; the third most frequent is WKKW, except again for 1.5 V, where it is WWKW. Hence, at most voltages, the most populated SF configurations sampled in the mutant are consistent with ion conduction occurring without water molecules intercalating ions, i.e., via the so-called hard or direct knock-on mechanism. At positive voltages, the filter remained in the conductive conformation, with only short-lived collapses (d1-min < 5 Å) when the site S2 is empty.

**Figure 4:**
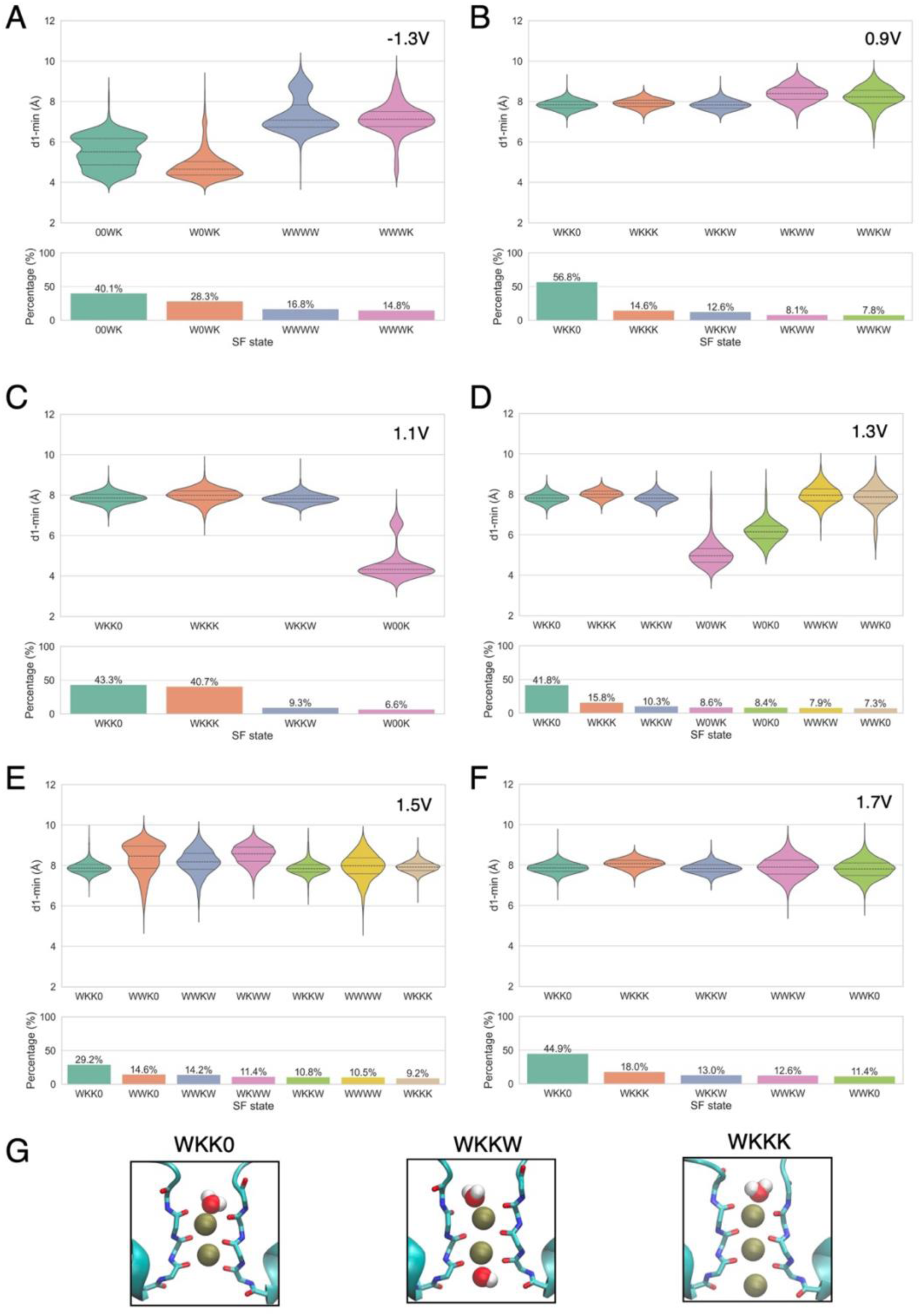
Configurations of ions in the A317T SF (partially open AG) during simulations with applied electric fields. (**A-F**) Top panels: violin plots of d1 values and ion conformations at different voltages; bottom panels: percentage occurrence of the configurations over all the trajectories. (**G**) 3D representations of the most common ion-water conformations in the SF.

**Table 3:** Summary of most and second most frequent ion-water filter conformations (partially open variant and closed WT)

|  |  | Voltage (V) |  |  |  |  |  |
| --- | --- | --- | --- | --- | --- | --- | --- |
|  |  | -1.3 | 0.9 | 1.1 | 1.3 | 1.5 | 1.7 |
| A317T | 1st | 00WK | WKK0 | WKK0 | WKK0 | WKK0 | WKK0 |
|  | 2nd | W0WK | WKKK | WKKK | WKKK | WWK0 | WKKK |
| WT | 1st | WKK0 | WKK0 | WKK0 | WKK0 | WKK0 | WKK0 |
|  | 2nd | WKWK | WKKK | WKK0 | WKKK | WKKK | WKKK |

In the WT system, while no complete permeation event was observed, the filter configurations can also change due to local fluctuations of the atoms. The most probable configuration is again, at all voltages, WKK0; the second most probable is WKKK, except at -1.3V and 1.1V, where it is WKWK and WKK0, respectively (Suppl. Fig. S23 and Table 3).

In conclusion, with the exception of the mutant at negative voltage, the most probable SF configuration (WKK0) is the same at all explored voltages in both the partially open mutant and the closed, non-conducting WT. The second most probable configuration is also predominantly shared (WKKK), with minor exceptions in both channels. This correspondence may result from the low frequency of complete permeation events observed for the variant in this configuration, which exhibits only limited gate opening and therefore supports reduced ion flux.

#### Open A317T and WT

To verify that, consistent with experimental observations (Nappi et al. 2024), the A317T substitution does not abolish channel function, we also simulated the open WT and mutant structures under applied transmembrane fields. During equilibration, the backbone RMSDs were stable around 2-3 Å for both systems (Suppl. Fig. S24). At all tested voltages, the structure of the SF is again preserved, as shown by the four coordination distances (d_s1– d_s4). Only d_s1 deviates from the cryo-EM value at the highest voltage, similarly to what was observed in the partially open/closed simulations, possibly because it is the most exposed one to the extracellular solution (Suppl. Fig. S25). Fig. 5 shows the cumulative count of K^+^ permeation events across the channel in the open structure, averaged over all replicas at each voltage, for both the A317T variant (Fig. 5A) and WT (Fig. 5B). Notably, the open mutant channel supports K^+^ permeation across all simulated voltages, consistent with its experimentally observed activity (Nappi et al. 2024). At variance with the equal unitary conductance estimated from noise analysis, however, we observe higher ion permeation in the variant than in the WT at each simulated voltage. Since the difference is more pronounced at higher voltages, it may reflect an artifact of the supraphysiological fields applied.

**Figure 5:**
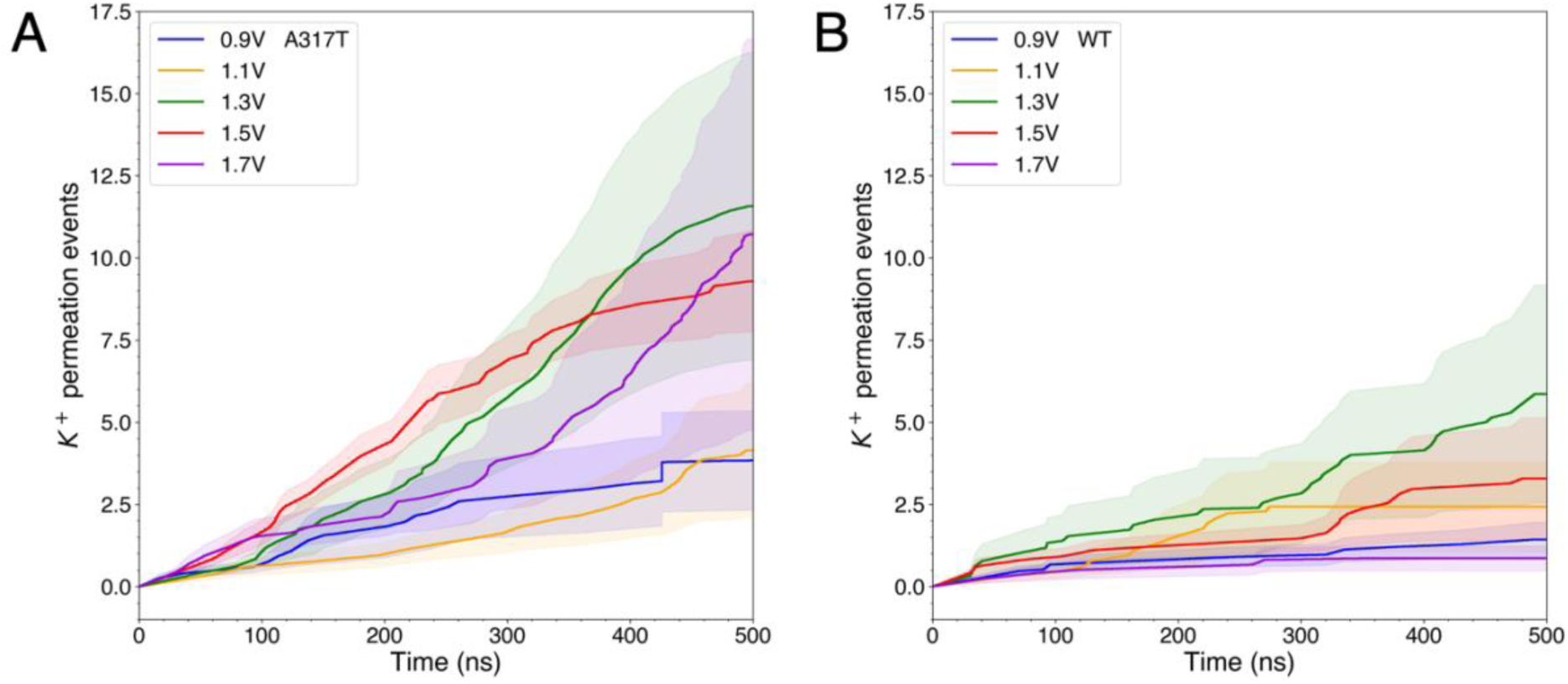
Time evolution of K^+^ permeation events through the open channel structures, averaged across replicas at each voltage. (**A**) A317T variant. (**B**) WT.

## Conclusions

GoF variants in the Kv7.2 channel are typically associated with severe neurodevelopmental and/or epileptic phenotypes. Elucidating the molecular mechanisms underlying variant-driven functional alterations, while potentially enabling therapeutic strategies, remains a challenging task. By providing an atomistic description of conformational states and interaction networks, MD simulations can help dissect how specific substitutions impact the channel functional properties.

Recently, we showed that three pathogenic variants located at the Kv7.2 AG produce GoF effects that were attributed to an increased probability of single-channel opening (Nappi et al. 2022, 2024). Our simulations of the mutants revealed that the closed AG conformation is destabilized by newly formed interactions that promote partial opening. Here, we sought to reinforce these findings by (i) extending the simulation timescales and (ii) showing that the variant-induced partial gate opening is sufficient to support ion permeation, consistent with electrophysiological recordings reporting measurable mutant currents at voltages where the WT channel remains closed or minimally conductive.

We also analyzed the impact of the force field on the emergence of the variant-driven, partially open gate conformation. CHARMM simulations of mutants starting from the closed AG, reaching up to 10 *μ*s for A317T, consistently sampled a conformation characterized by a wider gate and a hydrated cavity, indicating that the variants can favor transition of the closed state toward an open pore geometry. When simulated with AMBER, however, the mutants displayed more limited structural divergences from WT, with pores remaining closer to the native closed-state structure: still, while pore-radius profiles revealed minor differences, AG CDs captured more pronounced changes in the mutants. This comparison suggests that the extent of spontaneous gate widening depends, at least on the explored timescales, on the force field employed. This may reflect known differences among protein force-fields in backbone torsional preferences and secondary-structure stability, which can influence helix flexibility and thereby dampen or enhance the collective S_6_ motions required for Best et al. 2008; Lindorff-Larsen et al. 2010; Furini and Domene 2020). Indeed, previous simulations of K^+^ channels showed that CHARMM and AMBER descriptions can differ in their propensity to stabilize specific functional conformations and even in the resulting ion-permeation behavior (Ocello et al. 2020; Lam and Groot 2023).

Our applied-field simulations indicate that the CHARMM-derived partially open A317T conformation is functionally relevant, as it allows K permeation, whereas the corresponding WT closed structure remained non-conductive under the same conditions. Although these simulations used supraphysiological transmembrane potentials and restrained protein conformations, they were not intended to provide quantitative conductance estimates. Instead, they served as a functional test of pore geometry: even under a strong driving force, the WT closed structure remained non-conductive, whereas the partially open A317T conformation allowed K^+^ permeation. Conversely, both WT and A317T open structures displayed K^+^ translocation, suggesting that the variant primarily affects the closed-state AG architecture more than the conductive properties of the fully open pore.

Overall, this work provides structural insights into how GoF-associated variants can stabilize a partially open, yet conductive, intracellular gate in Kv7.2. By linking the pathological phenotype to specific structural rearrangements induced by mutations, these findings help identify a mechanistic framework for interpreting how clinically relevant KCNQ2 variants reshape channel behavior.

## Author Contributions

G.A. and L.M. designed the research. A.R. carried out all simulations and provided data curation. G.A. helped prepare the MD simulations. G.A., F.B., and L.M. supervised the project, provided resources, and provided project administration. F.M., M. T., F. B., L.M. provided funding acquisition. All authors analyzed the data and assisted in the investigation. G.A. and L.M. wrote the manuscript. All authors contributed to revising the article.

## Acknowledgments

We thank Pasquale Striano, Federico Zara, Mario Nappi, Carmen Domene, Simone Furini, and Anna Moroni for useful discussions. We acknowledge Alessandro Berselli for advice on MD analysis. We are grateful to Rossana Ciancio, Arta Mehilli, and Diego Moruzzo for administrative and technical assistance. We acknowledge ISCRA for awarding this project access to the LEONARDO supercomputer, owned by the EuroHPC Joint Undertaking and hosted by CINECA (Bologna, Italy), and the Data Science and Computation Facility and its Support Team for their support and assistance with the IIT High Performance Computing Infrastructure (Genova, Italy). This work was supported by the Italian Ministry for University and Research with PRIN2020 (project 2020XBFEMS to LM), PRIN2022 (project 2022M3KJ4N to MT), and PRIN2022 PNRR (project P2022FJXY5 to FM; project P2022ZANRF to MT); the European Union - Next Generation EU, Mission 4, Component 2, CUP E63C2 2002170007 (Project “A multiscale integrated approach to the study of the nervous system in health and disease”, MNESYS) (to MT); the Italian Ministry of Health with Ricerca Finalizzata (RF) Projects RF-2019-12370491 to MT and PNRR-MR1-2022-12376528 to FB and MT, and PNRR-MCNT2-2023-12377937 to MT; the European Partnership for Personalised Medicine EPPerMed 24_00187 (Validating in silico, in vitro, and in vivo BiomarkErs for personAlized Treatment in KCNQ2/3 encephalopathy (BEATKCNQ), the European Rare Disease Research Alliance (ERDERA) (VALidating the efficacy of a repUrposEd KCNQ opener in models of developmental and epileptic Encephalopathies, VALUEKCNQ); and by the Regione Campania NEURORARE Project (DGR 393 del 19/07/2022).

## References

Abraham, M. et al. 2023. “GROMACS 2023 Source Code.” 10.5281/zenodo.7588619.

Aksimentiev, A. et al. 2004. “Microscopic Kinetics of DNA Translocation Through Synthetic Nanopores.” J. Chem. Phys. 87: 2086–97.

Alberini, G. et al. 2021. “Structural Mechanism of ω-Currents in a Mutated Kv7.2 Voltage Sensor Domain from Molecular Dynamics Simulations.” J. Chem. Inf. Model. 61: 1354–67.

Balusek, C. et al. 2019. “Accelerating Membrane Simulations with Hydrogen Mass Repartitioning.” J. Chem. Theory Comput. 15: 4673–86.

Bekker, H. et al. 1993. “GROMACS—A Parallel Computer for Molecular-Dynamics Simulations.” In 4th International Conference on Computational Physics (PC 92), 252–56. World Scientific Publishing.

Berendsen, H. J. et al. 1995. “GROMACS: A Message-Passing Parallel Molecular Dynamics Implementation.” Comput. Phys. Commun. 91: 43–56.

Best, R. B. et al. 2008. “Are Current Molecular Dynamics Force Fields Too Helical?” Biophys. J. 95: L07–L09.

Best, R. B. et al. 2012. “Optimization of the Additive CHARMM All-Atom Protein Force Field Targeting Improved Sampling of the Backbone ϕ, ψ and Side-Chain χ1 and χ2 Dihedral Angles.” J. Chem. Theory Comput. 8: 3257–73.

Bouysset, C. et al. 2021. “ProLIF: A Library to Encode Molecular Interactions as Fingerprints.” J. Cheminform. 13: 72.

Crozier, P. S. et al. 2001. “Model Channel Ion Currents in NaCl-Extended Simple Point Charge Water Solution with Applied-Field Molecular Dynamics.” Biophys. J. 81: 3077–89.

Darden, T. et al. 1993. “Particle Mesh Ewald: An n Log (n) Method for Ewald Sums in Large Systems.” J. Chem. Phys. 98: 10089–99.

Daura, X. et al. 1999. “Peptide Folding: When Simulation Meets Experiment.” Angew. Chem. 38: 236–40.

Dickson, C. J. et al. 2022. “Lipid21: Complex Lipid Membrane Simulations with AMBER.” J. Chem. Theory Comput. 18: 1726–36.

Dirkx, N. et al. 2020. “The Role of Kv7.2 in Neurodevelopment: Insights and Gaps in Our Understanding.” Front. Physiol. 11.

Dunbrack, R. L., Jr. et al. 1993. “Backbone-Dependent Rotamer Library for Proteins Application to Side-Chain Prediction.” J. Mol. Biol. 230: 543–74.

Feenstra, K. A. et al. 1999. “Improving Efficiency of Large Time-Scale Molecular Dynamics Simulations of Hydrogen-Rich Systems.” J. Comput. Chem. 20: 786–98.

Feller, S. E. et al. 1995. “Constant Pressure Molecular Dynamics Simulation: The Langevin Piston Method.” J. Chem. Phys. 103: 4613–21.

Fiorin, G. et al. 2013. “Using Collective Variables to Drive Molecular Dynamics Simulations.” Mol. Phys. 111: 3345–62.

Furini, S. et al. 2020. “Critical Assessment of Common Force Fields for Molecular Dynamics Simulations of Potassium Channels.” J. Chem. Theory Comput. 16: 7148–59.

Goddard, T. D. et al. 2018. “UCSF ChimeraX: Meeting Modern Challenges in Visualization and Analysis.” Protein Sci. 27: 14–25.

Goto, A. et al. 2019. “Characteristics of KCNQ2 Variants Causing Either Benign Neonatal Epilepsy or Developmental and Epileptic Encephalopathy.” Epilepsia 60: 1870–80.

Gumbart, J. et al. 2012. “Constant Electric Field Simulations of the Membrane Potential Illustrated with Simple Systems.” Biochim. Biophys. Acta 1818: 294–302.

Hess, B. et al. 1997. “LINCS: A Linear Constraint Solver for Molecular Simulations.” J. Comput. Chem. 18: 1463–72.

Hoover, W. 1985. “Canonical Dynamics: Equilibrium Phase-Space Distributions.” Phys. Rev. A 31: 1695.

Hopkins, C. W. et al. 2015. “Long-Time-Step Molecular Dynamics Through Hydrogen Mass Repartitioning.” J. Chem. Theory Comput. 11: 1864–74.

Hoshi, N. 2020. “M-Current Suppression, Seizures and Lipid Metabolism: A Potential Link Between Neuronal Kv7 Channel Regulation and Dietary Therapies for Epilepsy.” Front. Physiol. 11.

Huang, J. et al. 2017. “CHARMM36m: An Improved Force Field for Folded and Intrinsically Disordered Proteins.” Nat. Methods 14: 71–73.

Huang, J. et al. 2013. “CHARMM36 All-Atom Additive Protein Force Field: Validation Based on Comparison to NMR Data.” J. Comput. Chem. 34: 2135–45.

Humphrey, W. et al. 1996. “VMD: Visual Molecular Dynamics.” J. Mol. Graph. 14: 33–38.

Jepps, T. A. et al. 2021. “Editorial. Kv7 Channels: Structure, Physiology, and Pharmacology.” Front. Physiol. 12.

Jo, S. et al. 2007. “Automated Builder and Database of Protein/Membrane Complexes for Molecular Dynamics Simulations.” PloS One 2: e880.

Jorgensen, W. L. et al. 1983. “Comparison of Simple Potential Functions for Simulating Liquid Water.” J. Chem. Phys. 79: 926–35.

Joung, I. S. et al. 2008. “Determination of Alkali and Halide Monovalent Ion Parameters for Use in Explicitly Solvated Biomolecular Simulations.” J. Phys. Chem. B 112: 9020–41.

Khalili-Araghi, F. et al. 2006. “Dynamics of K+ Ion Conduction Through Kv1.2.” Biophys. J. 91: L72–74.

Klauda, J. B. et al. 2010. “Update of the CHARMM All-Atom Additive Force Field for Lipids: Validation on Six Lipid Types.” J. Phys. Chem. B 114: 7830–43.

Kopec, W. et al. 2019. “Molecular Mechanism of a Potassium Channel Gating Through Activation Gate-Selectivity Filter Coupling.” Nat. Commun. 10.

Kutzner, C. et al. 2011. “Computational Electrophysiology: The Molecular Dynamics of Ion Channel Permeation and Selectivity in Atomistic Detail.” Biophys. J. 101: 809–17.

Lam, C. K. et al. 2023. “Ion Conduction Mechanisms in Potassium Channels Revealed by Permeation Cycles.” J. Chem. Theory Comput. 19: 2574–89.

Li, X. et al. 2021. “Molecular Basis for Ligand Activation of the Human KCNQ2 Channel.” Cell Res. 31: 52–61.

Lindorff-Larsen, K. et al. 2010. “Improved Side-Chain Torsion Potentials for the Amber ff99SB Protein Force Field.” Proteins 78: 1950–58.

Lomize, M. A. et al. 2012. “OPM Database and PPM Web Server: Resources for Positioning of Proteins in Membranes.” Nucleic Acids Res. 40: D370–76.

Luo, Y. et al. 2010. “Simulation of Osmotic Pressure in Concentrated Aqueous Salt Solutions.” J. Phys. Chem. Lett. 1: 183–89.

Ma, D. et al. 2023. “Ligand Activation Mechanisms of Human KCNQ2 Channel.” Nat. Commun. 14: 6632.

Machtens, J.-P. et al. 2015. “Mechanisms of Anion Conduction by Coupled Glutamate Transporters.” Cell 160: 542–53.

Maier, J. A. et al. 2015. “ff14SB: Improving the Accuracy of Protein Side Chain and Backbone Parameters from ff99SB.” J. Chem. Theory Comput. 11: 3696–713.

Malerba, F. et al. 2020. “Genotype-Phenotype Correlations in Patients with de Novo KCNQ2 Pathogenic Variants.” Neurol. Genet. 6: e528.

Martyna, G. J. et al. 1994. “Constant Pressure Molecular Dynamics Algorithms.” J. Chem. Phys. 101: 4177–89.

Meng, E. C. et al. 2023. “UCSF ChimeraX: Tools for Structure Building and Analysis.” Protein Sci. 32: e4792.

Miceli, F. et al. 2015. “Early-Onset Epileptic Encephalopathy Caused by Gain-of-Function Mutations in the Voltage Sensor of Kv7.2 and Kv7.3 Potassium Channel Subunits.” J. Neurosci. 35: 3782–93.

Miceli, F. et al. 2022. “KCNQ2 R144 Variants Cause Neurodevelopmental Disability with Language Impairment and Autistic Features Without Neonatal Seizures Through a Gain-of-Function Mechanism.” EBioMedicine 81: 104130.

Michaud-Agrawal, N. et al. 2011. “MDAnalysis: A Toolkit for the Analysis of Molecular Dynamics Simulations.” J. Comput. Chem. 32: 2319–27.

Millichap, J. J. et al. 2017. “Infantile Spasms and Encephalopathy Without Preceding Neonatal Seizures Caused by KCNQ2 R198Q, a Gain-of-Function Variant.” Epilepsia 58: e10–e15.

Miyamoto, S. et al. 1992. “Settle: An Analytical Version of the SHAKE and RATTLE Algorithm for Rigid Water Models.” J. Comput. Chem. 13: 952–62.

Nappi, P. et al. 2020. “Epileptic Channelopathies Caused by Neuronal Kv7 (KCNQ) Channel Dysfunction.” Pflugers Arch. 472: 881–98.

Nappi, M. et al. 2024. “Constitutive Opening of the Kv7.2 Pore Activation Gate Causes KCNQ2-Developmental Encephalopathy.” Proc. Natl. Acad. Sci. USA 121: e2412388121.

Nappi, M. et al. 2022. “Gain of Function Due to Increased Opening Probability by Two KCNQ5 Pore Variants Causing Developmental and Epileptic Encephalopathy.” Proc. Natl. Acad. Sci. USA 119: e2116887119.

Nosé, S. 1984. “A Molecular Dynamics Method for Simulations in the Canonical Ensemble.” Mol. Phys. 52: 255–68.

Noskov, S. Y. et al. 2008. “Control of Ion Selectivity in LeuT: Two Na+ Binding Sites with Two Different Mechanisms.” J. Mol. Biol. 377: 804–18.

Ocello, R. et al. 2020. “Conduction and Gating Properties of the TRAAK Channel from Molecular Dynamics Simulations with Different Force Fields.” J. Chem. Inf. Model. 60: 6532– 43.

Parrinello, M. et al. 1981. “Polymorphic Transitions in Single Crystals: A New Molecular Dynamics Method.” J. Appl. Phys. 52: 7182–90.

Pettersen, E. F. et al. 2004. “UCSF Chimera—a Visualization System for Exploratory Research and Analysis.” J. Comput. Chem. 25: 1605–12.

Pettersen, E. F. et al. 2021. “UCSF ChimeraX: Structure Visualization for Researchers, Educators, and Developers.” Protein Sci. 30: 70–82.

Phillips, J. C. et al. 2005. “Scalable Molecular Dynamics with NAMD.” J. Comput. Chem. 26: 1781–802.

Ryckaert, J.-P. et al. 1977. “Numerical Integration of the Cartesian Equations of Motion of a System with Constraints: Molecular Dynamics of n-Alkanes.” J. Comput. Phys. 23: 327–41.

Smart, O. S. et al. 1997. “A Novel Method for Structure-Based Prediction of Ion Channel Conductance Properties.” Biophys. J. 72: 1109.

Smart, O. S. et al. 1996. “HOLE: A Program for the Analysis of the Pore Dimensions of Ion Channel Structural Models.” J. Mol. Graph. 14: 354–60.

Sotomayor, M. et al. 2007. “Ion Conduction Through MscS as Determined by Electrophysiology and Simulation.” Biophys. J. 92: 886–902.

Steinbach, P. J. et al. 1994. “New Spherical-Cutoff Methods for Long-Range Forces in Macromolecular Simulation.” J. Comput. Chem. 15: 667–83.

Sun, J. et al. 2020. “Structural Basis of Human KCNQ1 Modulation and Gating.” Cell 180: 340–47.

Venable, R. M. et al. 2013. “Simulations of Anionic Lipid Membranes: Development of Interaction-Specific Ion Parameters and Validation Using NMR Data.” J. Phys. Chem. B 117: 10183–92.

Weckhuysen, S. et al. 2012. “KCNQ2 Encephalopathy: Emerging Phenotype of a Neonatal Epileptic Encephalopathy.” Ann. Neurol. 71: 15–25.

Zuberi, S. M. et al. 2022. “ILAE Classification and Definition of Epilepsy Syndromes with Onset in Neonates and Infants: Position Statement by the ILAE Task Force on Nosology and Definitions.” Epilepsia 63: 1349–97.

