## Supplementary file for "Molecular mechanism of partially open Kv7.2 gate stabilization by gain-of-function variants"

### S1 Methods for Constant electric field simulations

#### S1.1 Simulations set up

### Nonequilibrium molecular dynamics simulations with the electric field. Equilibration Protocol

Before applying the electric field, systems were equilibrated via a two-stage protocol (NPT followed by NVT), with different procedures for the initially closed- (Fig. S1) open-gate conformations, summarized hereafter and in Figs S2 and S3, respectively.

#### Preliminary equilibration for the starting closed AG conformation

NPT stage:

1. C-GUI standard equilibration protocol (from C-GUI step 6.1 to step 6.6), followed by additional 60 ns of extended step 6.6.
2. 10 ns with standard atomic masses and a time step of $\Delta t$=2 fs. D Harmonic potential dihedral restraints were applied to the SF (from residue 276 to residue 283) to prevent its collapse, with a force constant of 50 kcal/mol.
3. 1000 ns with HMR and a time step of $\Delta t$=4 fs. The same SF restraints of step 2 here above were applied.
4. 250 ns with standard atomic masses and a time step $\Delta t$=2 fs. The same SF restraints of step 2 and 3 here above were applied.

To identify a a representative structure for the following NVT steps, we performed a cluster analysis on the last 250 ns using the gromos algorithm with a cutoff of 1.5 Å. The selected structure is characterized by the following features, based on our previous MD simulations (Nappi, Alberini et al., 2024 Proc Natl Acad Sci USA. 121:e2412388121):

- partially open AG (d2 = 12.3 ± 0.9 Å; d3 = 14.81 ± 1.04 Å);
- hydrated PD (more than 10 water molecules in the central cavity, residues G313 to G279);
- open SF (d1 = 8.66 ± 1.06 Å).

The selected structure was then used as starting conformation for the following five equilibration steps in NVT:

1. 10 ns with standard atomic masses and a time step of $\Delta t$=1 fs, positional restraints on all protein heavy atoms.

2. 10 ns with standard atomic masses and a time step of $\Delta t$=2 fs, positional restraints on the protein backbone.

3. 10 ns with standard atomic masses and a time step of $\Delta t$=2 fs, positional restraints on protein C$\alpha$ atoms.

4. 10 ns with standard atomic masses and a time step of $\Delta t$=2 fs, positional restraints on TM C$\alpha$ residues (residue 238 to 247 and 294 to 303 of each helix).

5. 10 ns with standard atomic masses and a time step of $\Delta t$=2 fs, positional restraints on C$\alpha$ of the TM residues, and lower-bound restraints on the inter-subunit distances between C$\alpha$ atoms of C-terminal residues (from 325 to 330) to move them away from the AG (flat-bottom, half-harmonic restraining potentials with threshold value -LowerWall keyword- 35 Å, and force costant of 100 kcal/mol). These restraints were used because, after removal of the calmodulin domains, the protein C-termini were free to fluctuate and could potentially occlude the AG, creating a barrier to ion entry.

6. Gradual application of the external electric field. 10ns MD simulation per run, 200mV increments, starting from 100mV, until the desired value is reached.

#### Preliminary equilibration for the open AG conformation

For the open AG conformation, the NPT phase was modified to preserve gate hydration and geometry.

NPT phase (2 steps):

1. C-GUI standard equilibration protocol, followed by additional 60 ns of extended step 6.6.

2. 500 ns with standard masses (time step $\Delta t$ = 2 fs), with lower-bound restraints on AG cross distances d2 and d3 to allow proper hydration of the channel pore. A LowerWall of 13.50 Åwas set for d2, while for d3 it was 19.60 Å. The force costant was 100 kcal/mol/Å$^2$ in both cases.

For the subsequent NVT phase, we followed the same procedure described above for the initially closed AG structures. In this case, d2 was maintained between 15.04 Å and 15.76 Å for the WT and between 14.92 Å and 15.92 Å for A317T. Similarly, d3 was maintained between 18.05 Å and 19.29 Å for WT and between 18.23 Å and 19.65 Å for A317T.

#### S2 Supplementary figures

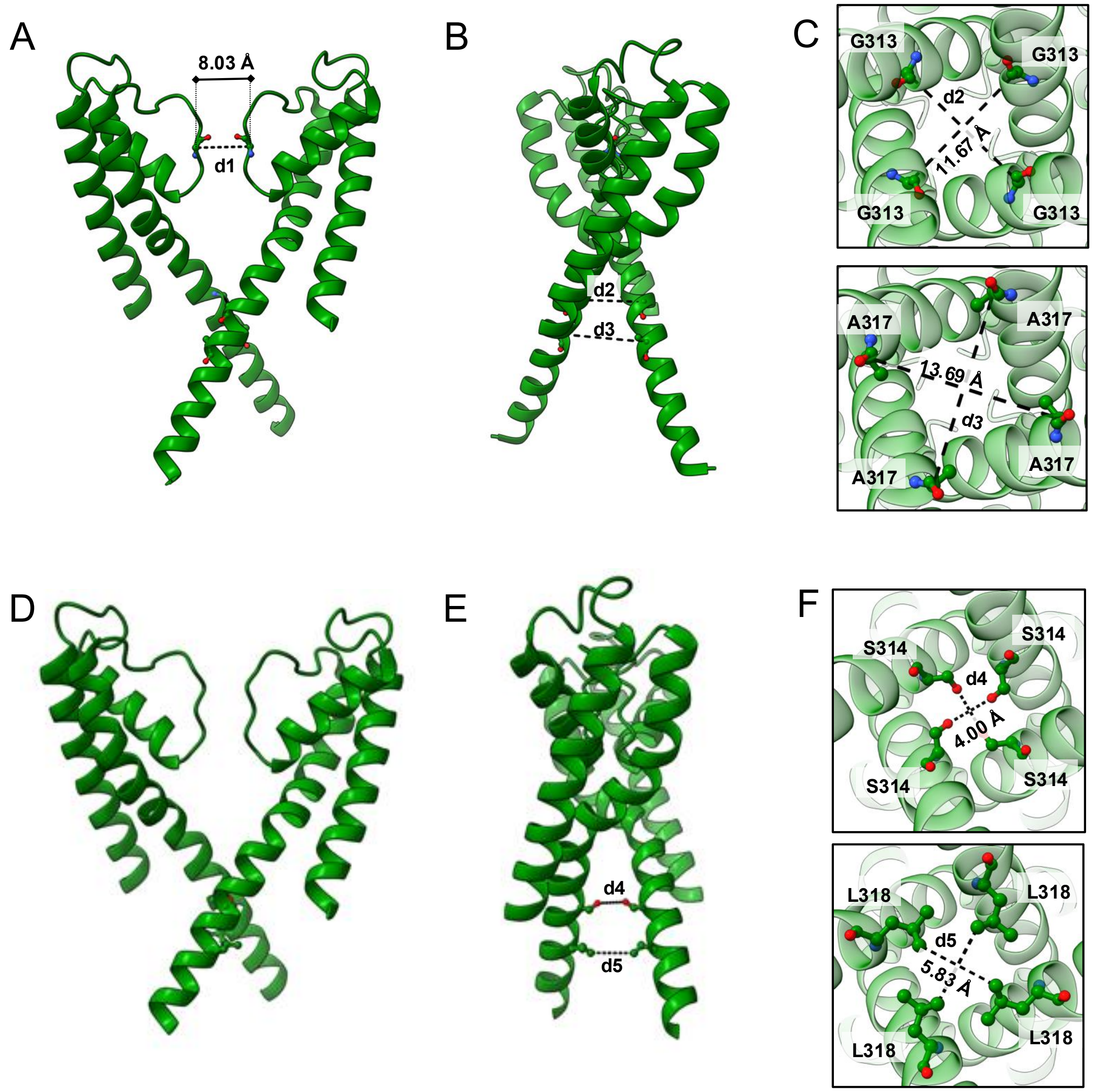


Figure S1: Definition of cross distances (CDs) and their illustration and values in the WT Kv7.2 cryo-EM structure with a closed AG (PDB ID: 7CR0). (**A**) d1, between diagonally opposed G279 C$\alpha$ atoms (8.03 Å in the closed AG structure). (**B**) d2, between diagonally opposed G313 C$\alpha$ (11.67 Å in the closed AG structure). (**C**) d3, between diagonally opposed A317 C$\alpha$ atoms (13.69 Å in the closed AG structure). (**D** -**F**) d4 and d5, between S314 O$\gamma$ atoms and L318 C$\delta$ atoms, respectively (4 Å and 5.83 Å in the closed AG structure).

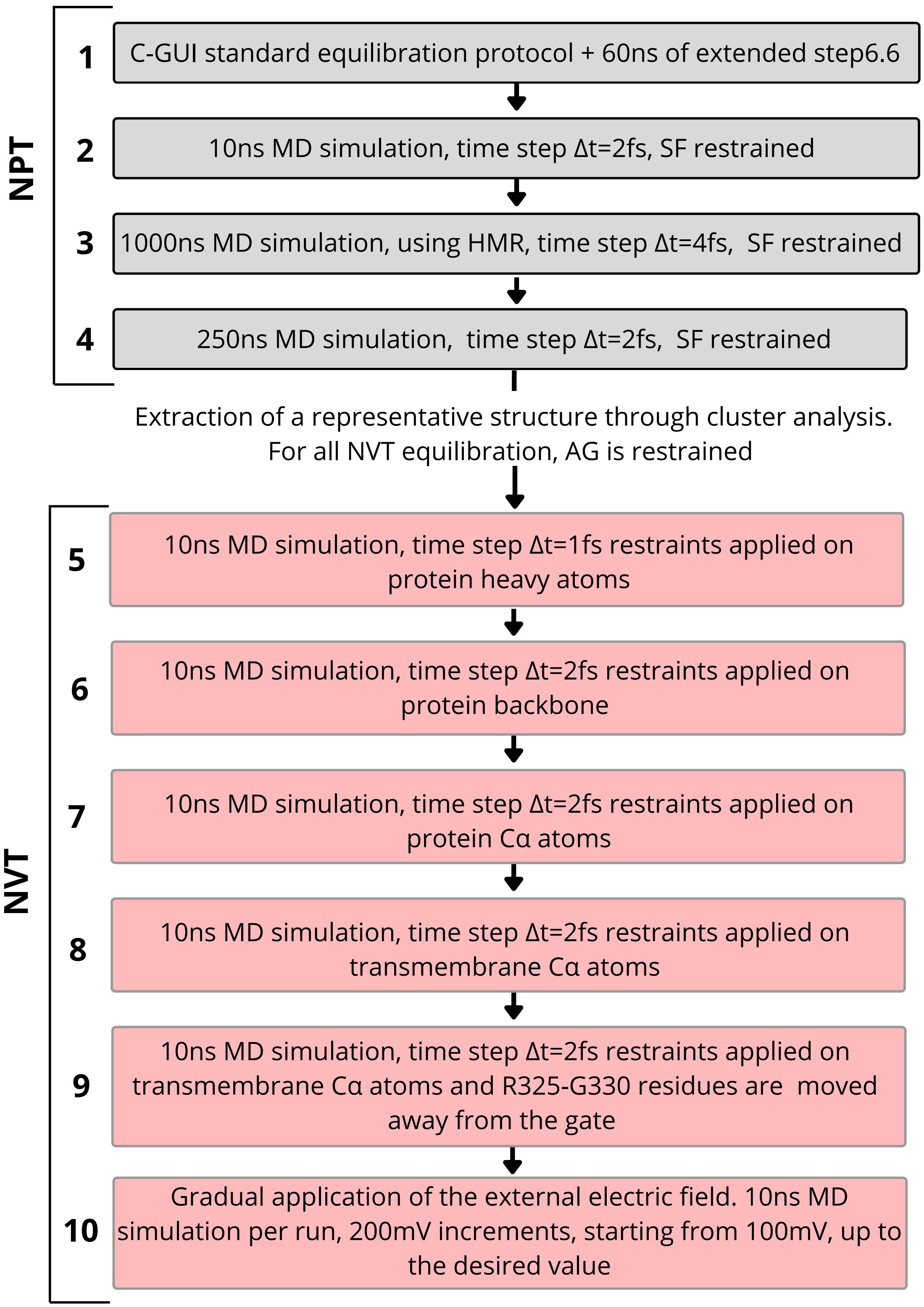


Figure S2: Equilibration protocol - Initially closed States.

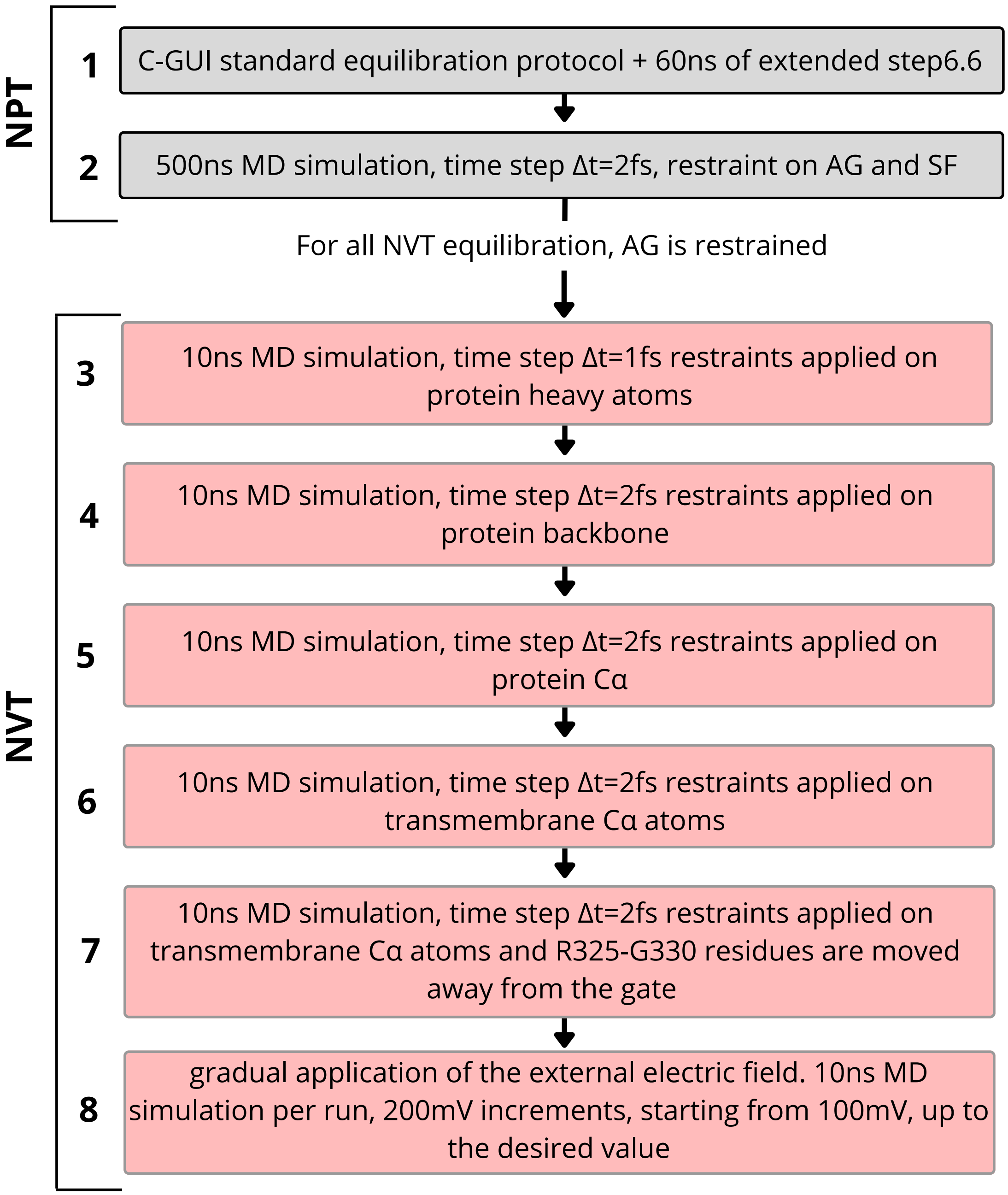


Figure S3: Equilibration protocol - Open States.

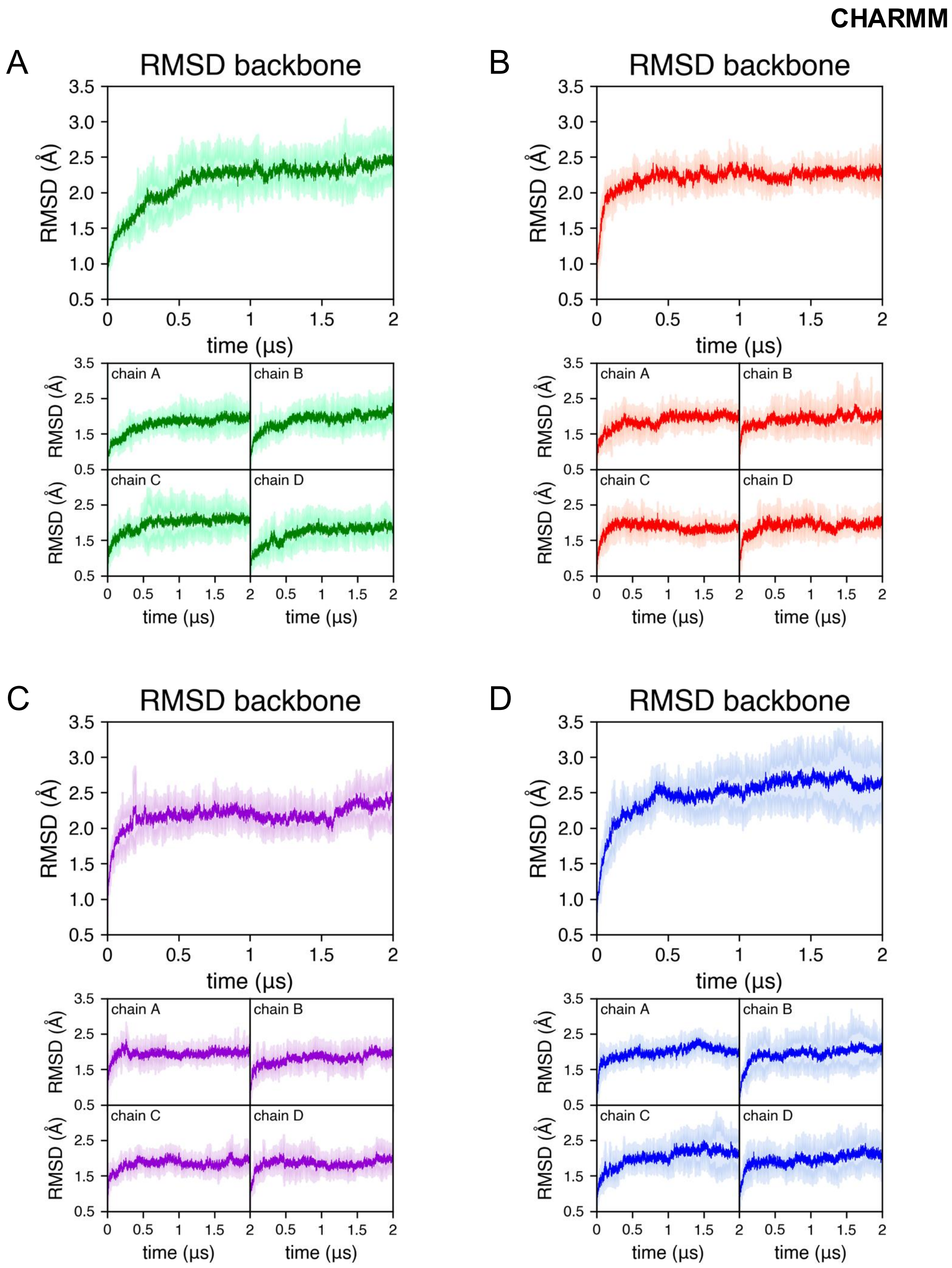


Figure S4: RMSD of the TM backbones in standard, equilibrium MD simulations using CHARMM36m, closed AG starting structures. RRMSDs were averaged over all five replicas of each channel: (**A**) WT (green), (**B**) G313S (red), (**C**) A317T (purple), (**D**) L318V (blue). Shaded areas represent standard deviations. In each panel, the bottom plots report values for different channel subunits.

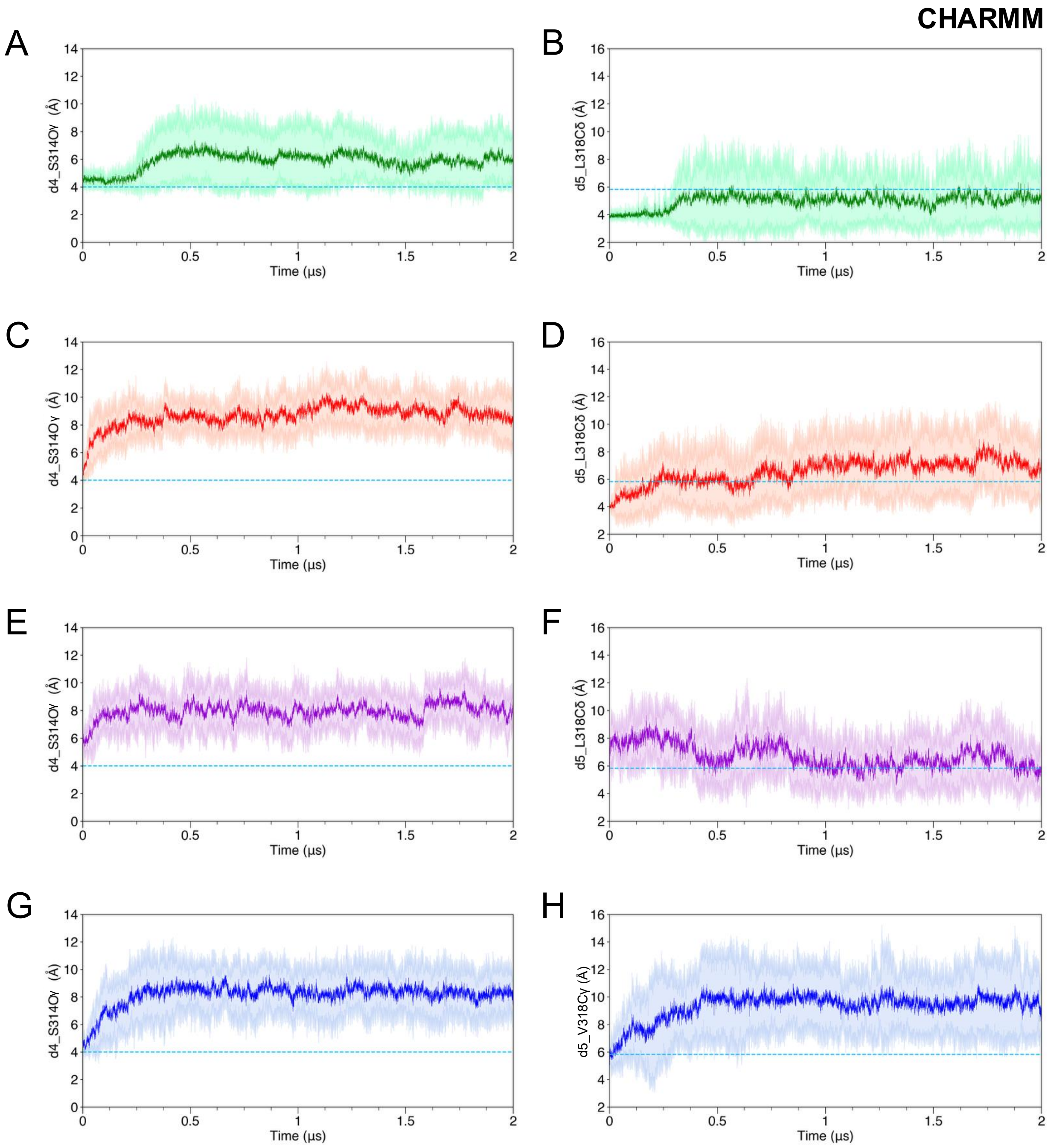


Figure S5: AG cross-distances during standard equilibrium simulations using CHARMM36m, closed AG starting structures. **(A, C, E, G)** Distance d4 for the four channels (green, WT; red, G313S; purple, A317T; blue, L318V) averaged over replicas (shaded areas represent standard deviations). **(B, D, F, H)** Distance d5 for the four channels, averaged over replicas. The dashed lines represent reference values in the closed AG structure.

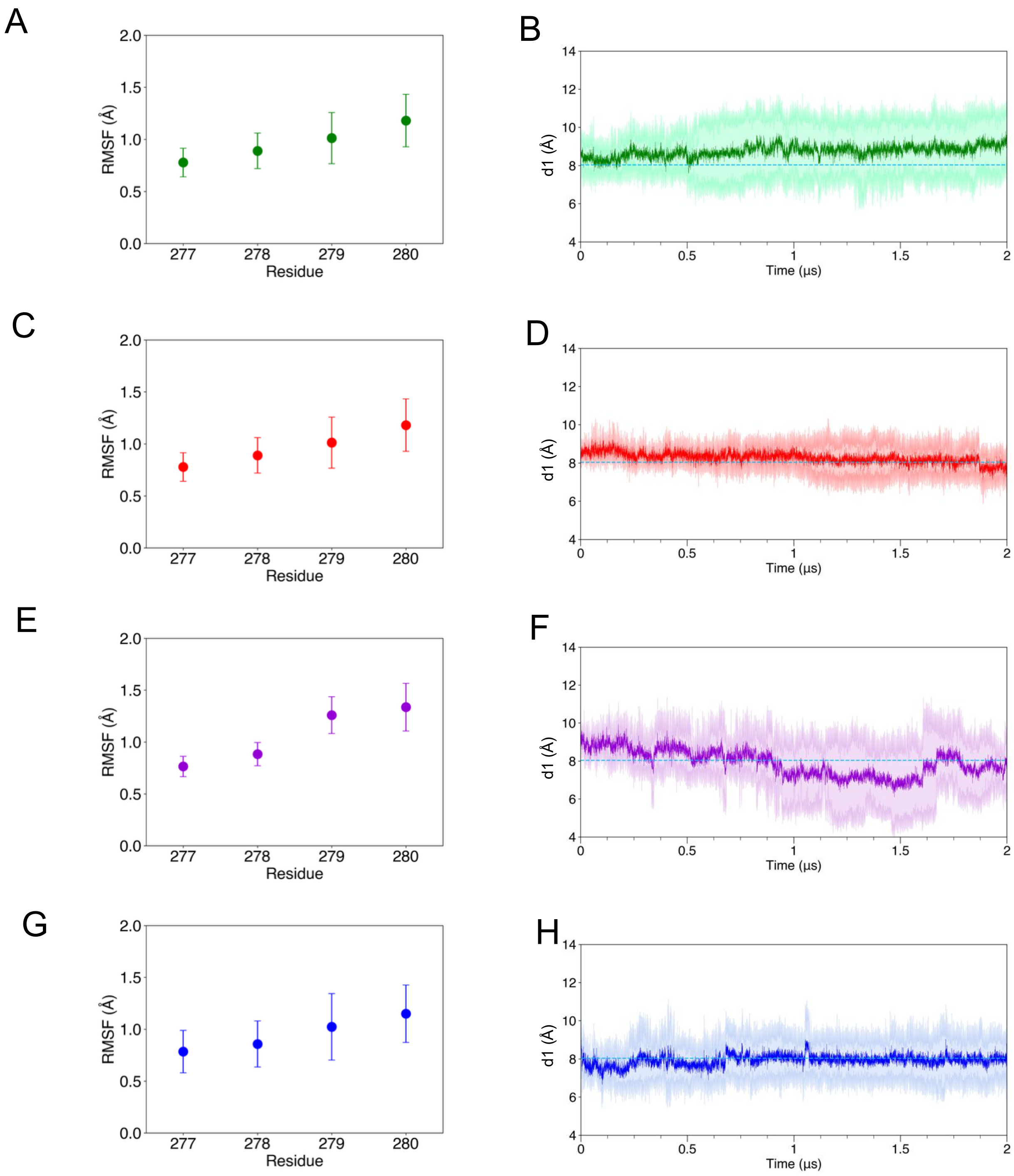


Figure S6: SF structural stability during simulations using Charmm36m, closed AG starting structures. (**A**, **C**, **E**, **G**) RMSF values for SF backbone atoms (residues T277 to Y280), averaged over all replicas. (**B**, **D**, **F**, **H**) CD d1 averaged over all replicas (shaded areas represent standard deviations).

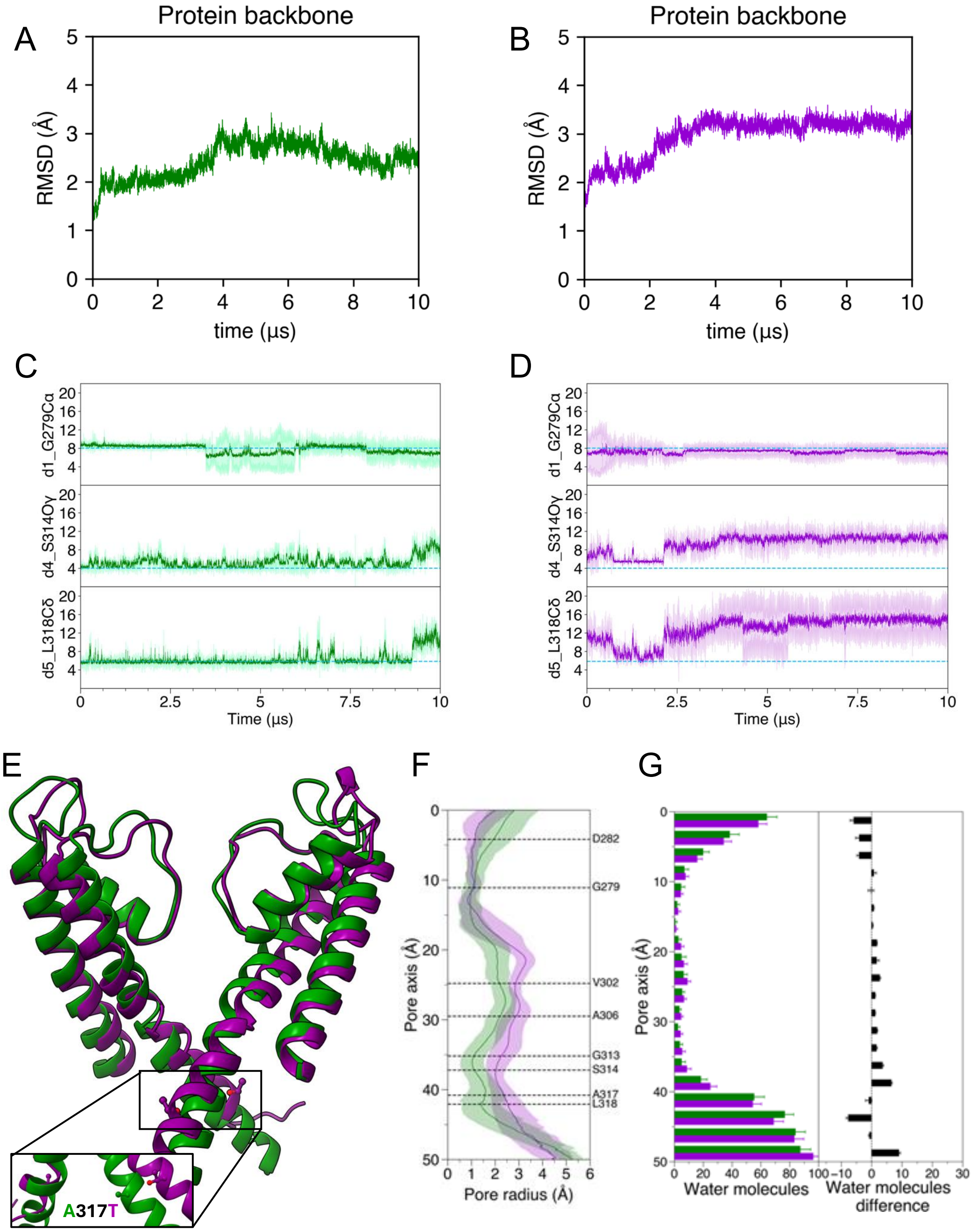


Figure S7: Analysis of 10 $\mu$s-long standard MD simulations using CHARMM36m, closed AG starting structures. (**A**, **B**) Backbone TM RMSDs (WT, green, and A317T, purple). (**C**, **D**) CDs d1, d4 and d5, for WT (green) and A317T (purple). Solid lines are averages of the two equivalent distances measured between atom pairs in opposing subunits (shaded areas represent standard deviations). The dashed lines are reference values in the cryo-EM structure (d1: 8.03 Å, d4: 4.00 Å, d5: 5.83 Å). (**E**) Superposition of the WT (green) and A317T (purple) structures at the end of the trajectories. (**F**) Channel radius along the pore axis averaged over all replicas (shaded areas are standard deviations). (**G**) Distribution of water molecules along the channel axis. Values are averages from all replicas.

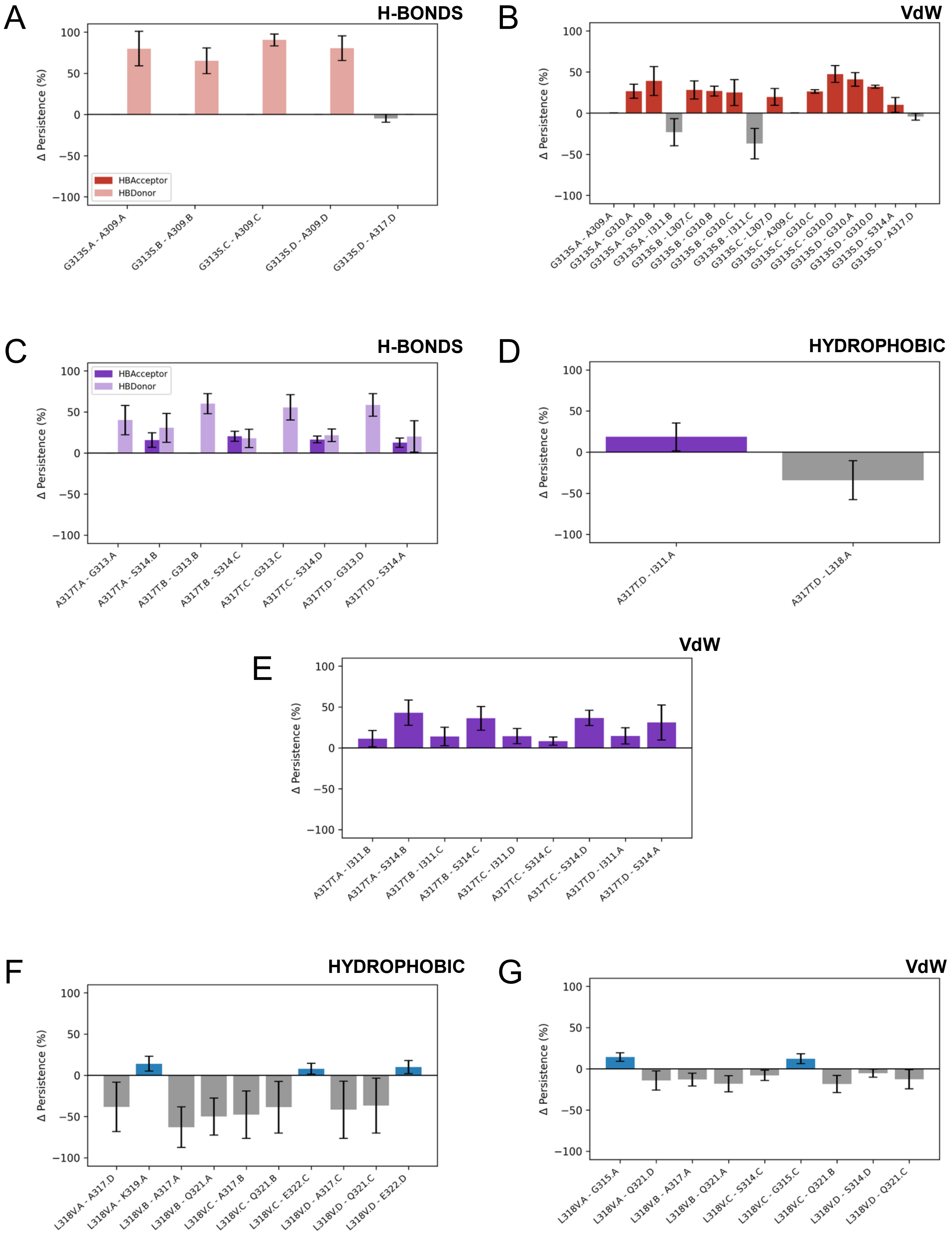


Figure S8: Differences in interaction persistence in CHARMM36m MD simulations, shown as mutant minus WT. (**A-B**) variant G313S, (**C-E**) variant A31T7, (**F-G**) variant L318V.

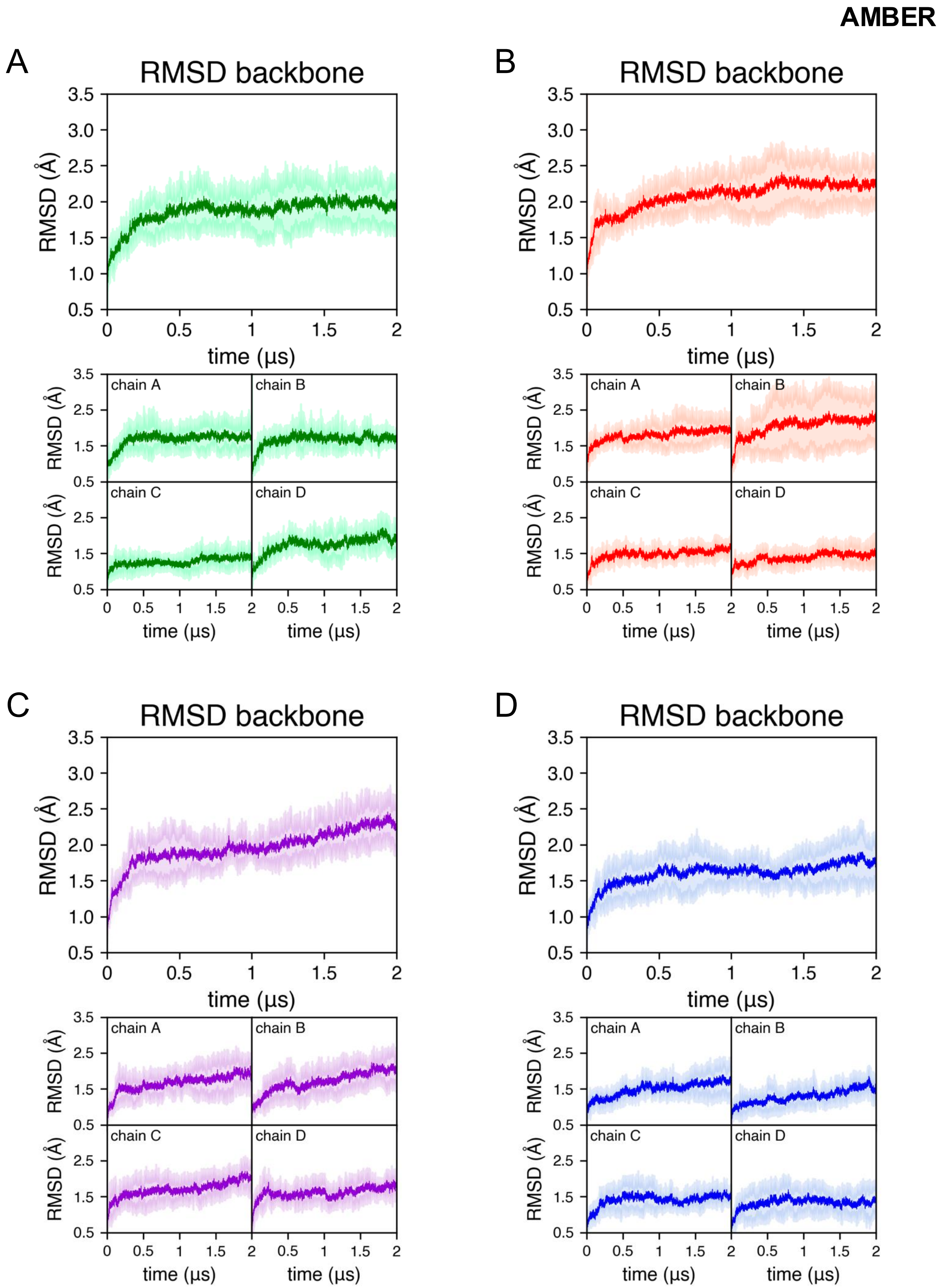


Figure S9: RMSD of the TM backbones in standard, equilibrium MD simulations using AMBER14SB, closed AG starting structures. RMSDs were averaged over all five replicas of each channel: (**A**) WT (green), (**B**) G313S (red), (**C**) A317T (purple), (**D**) L318V (blue). Shaded areas represent standard deviations. In each panel, the bottom plots report values for different channel subunits.

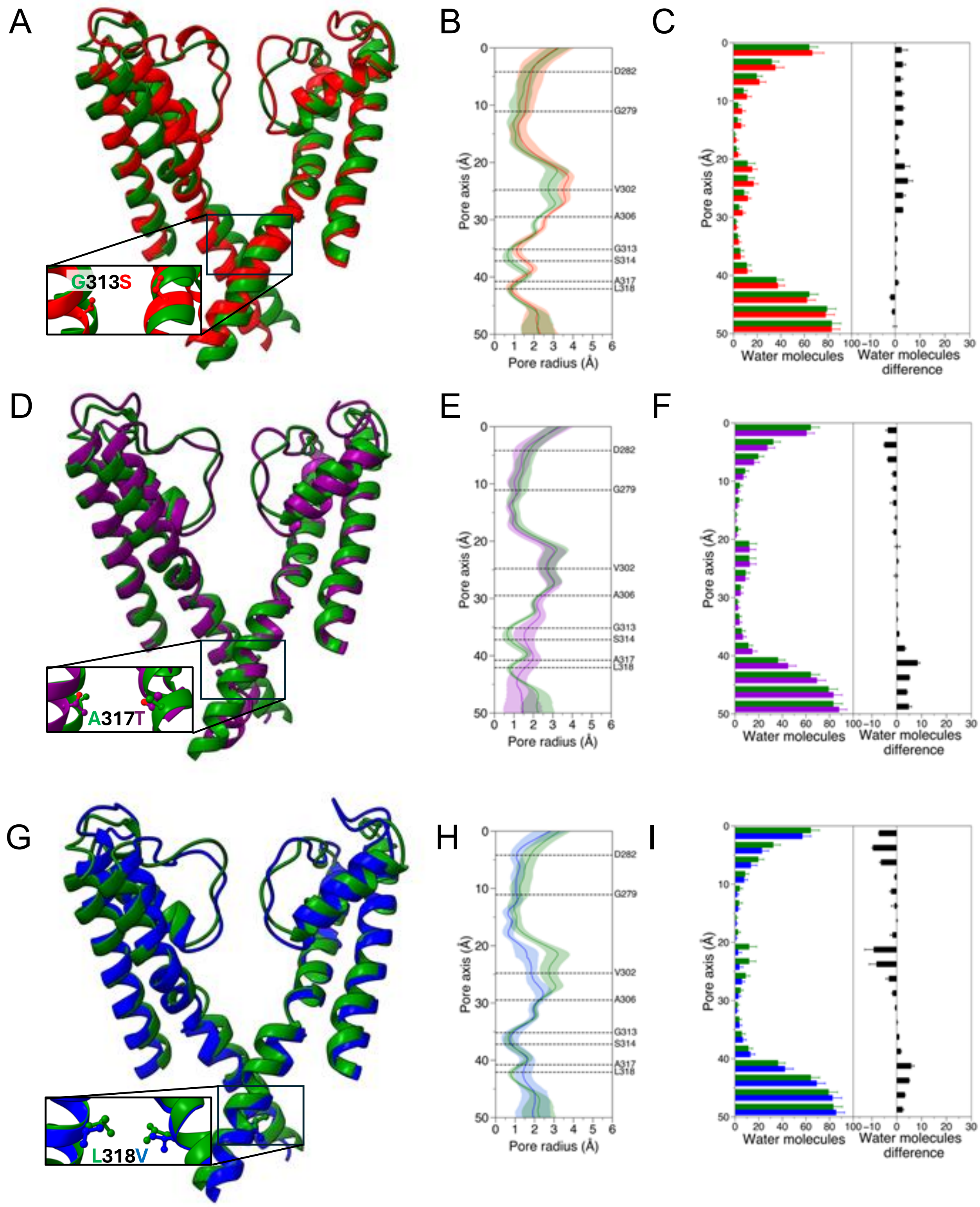


Figure S10: AMBER14SB MD simulations of WT and mutated channels, starting from the closed AG. (**A, D, G**) Superposition of WT (green) and variant structures at the end of 2 $\mu$s trajectories: G313S (red), A317T (purple), and L318V (blue). (**B, E, H**) Channel pore radius profiles averaged over all simulated replicas. Shaded regions represent the associated standard deviations. (**C, F, I**) Distribution of water molecules within the channel pores and differences between mutant and WT (panels on the right). Averages and standard deviations are calculated using all replicas

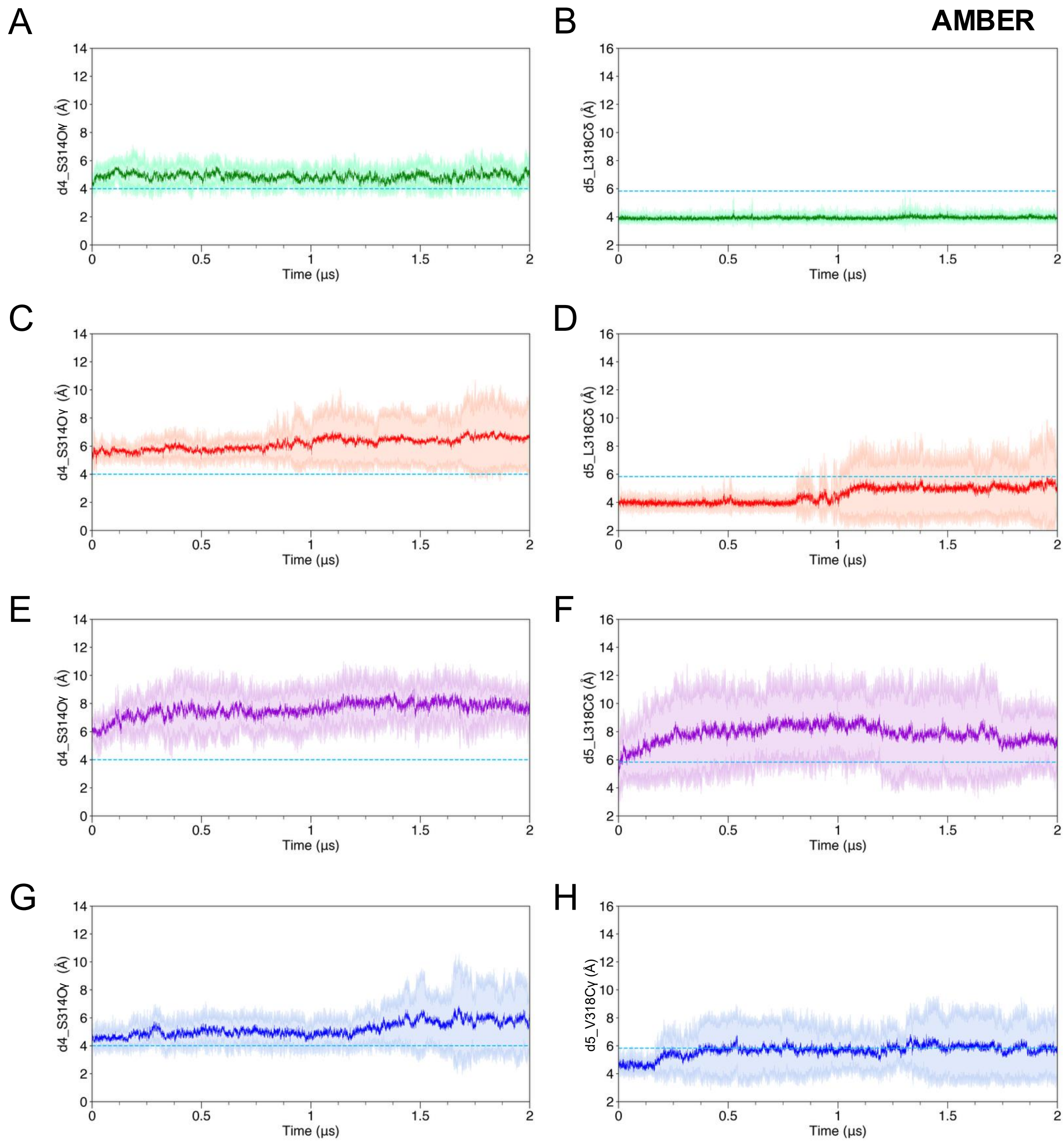


Figure S11: AG cross-distances during standard equilibrium simulations using AMBER14SB, closed AG starting structures. **(A, C, E, G)** Distance d4 for the four channels, averaged over replicas (shaded areas are standard deviations). The dashed line represents the reference value in the closed AG structure (PDB ID: 7CR0). **(B, D, F, H)** Distance d5 for the four channels, averaged over replicas (shaded areas are standard deviations). The dashed line represents the reference value in the closed AG structure.

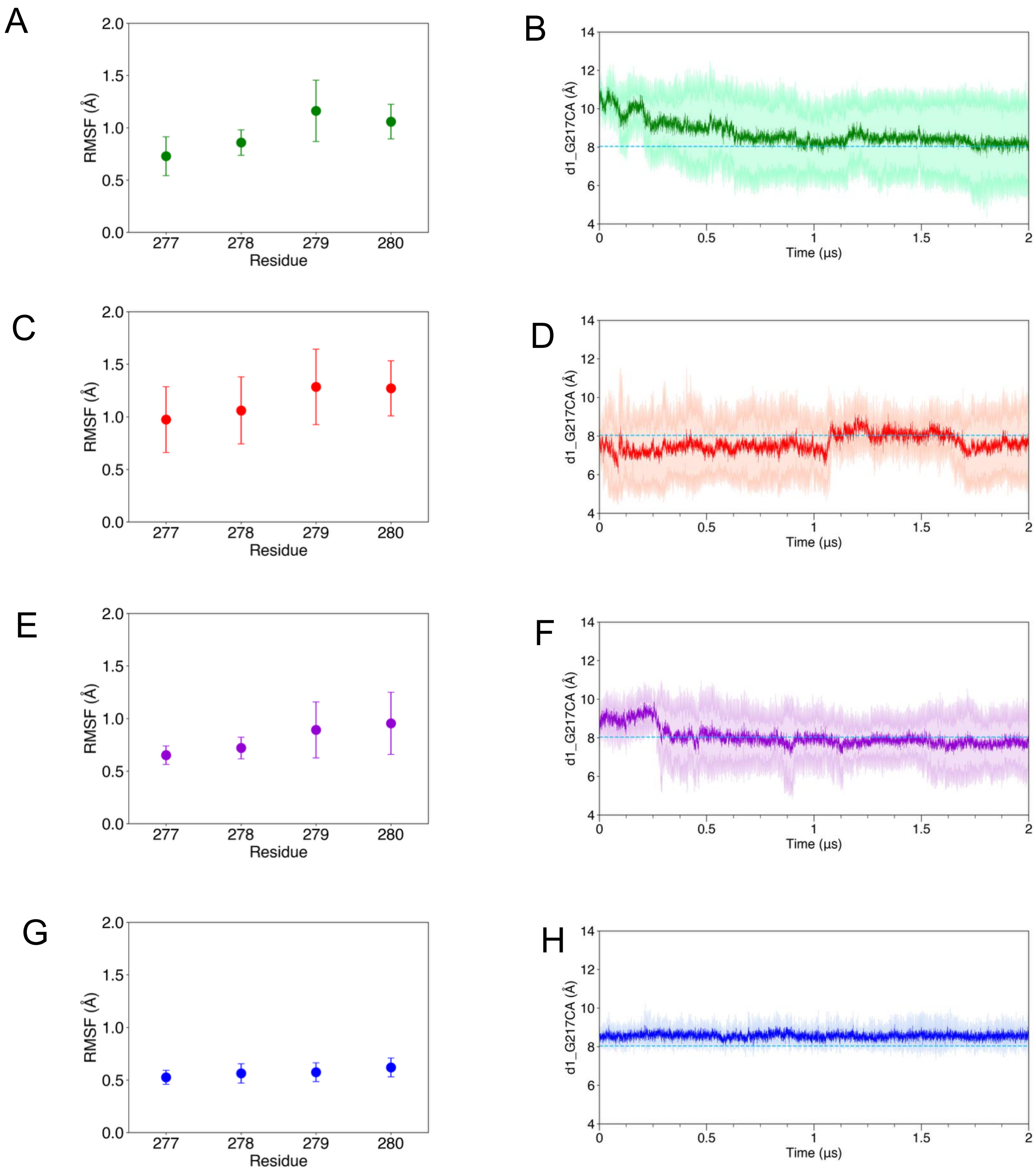


Figure S12: SF structural stability during simulations using AMBER14SB, closed AG starting structures. (**A, C, E, G**) RMSF values for SF backbone atoms (residues T277 to Y280), averaged over all replicas. (**B, D, F, H**) CD d1 averaged over all replicas (shaded areas are standard deviations).

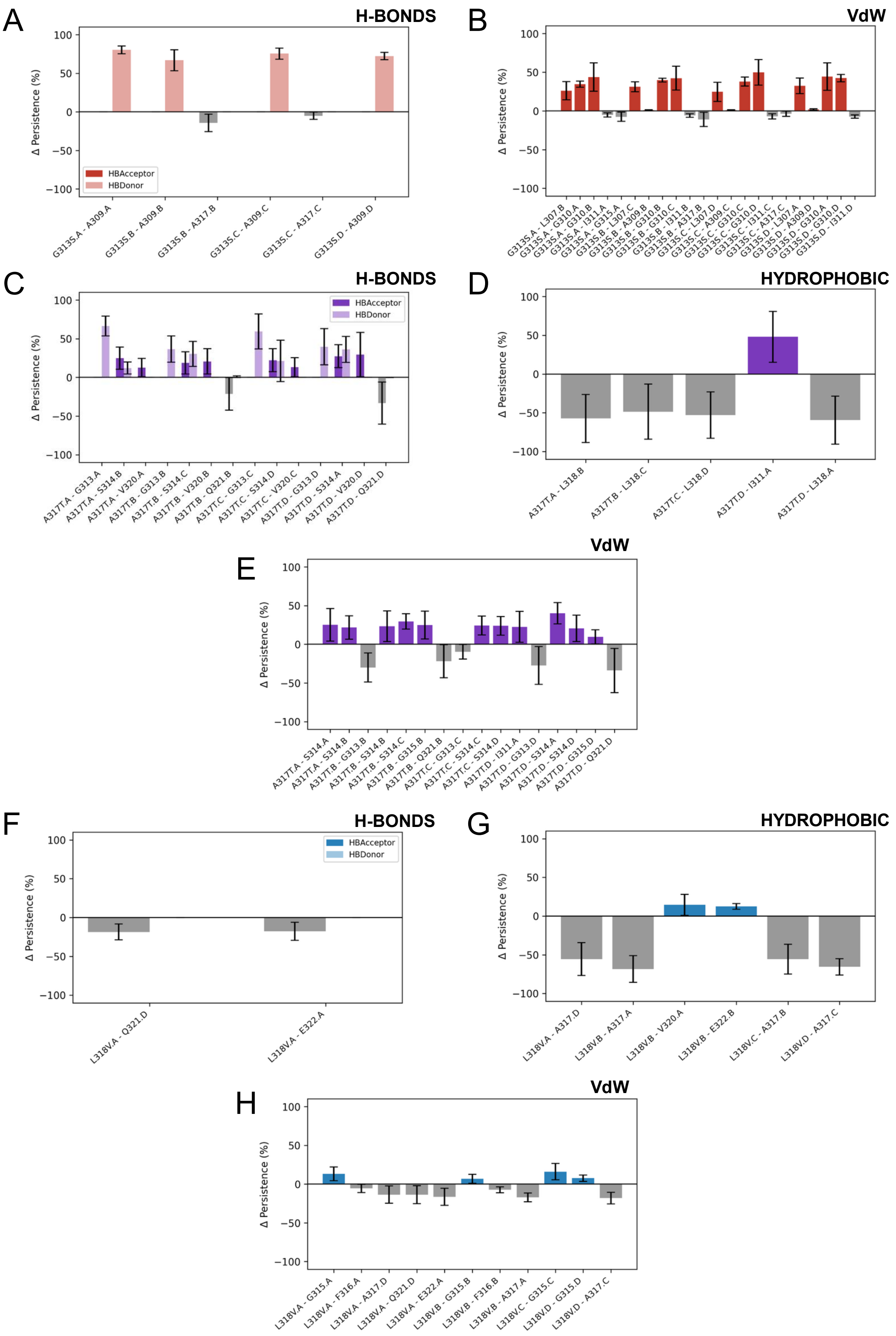


Figure S13: Differences in interaction persistence in AMBER14SB MD simulations, shown as mutant minus WT. (**A,B**) variant G313S, (**C-E**) variant A31T7, (**F,G**) variant L318V.

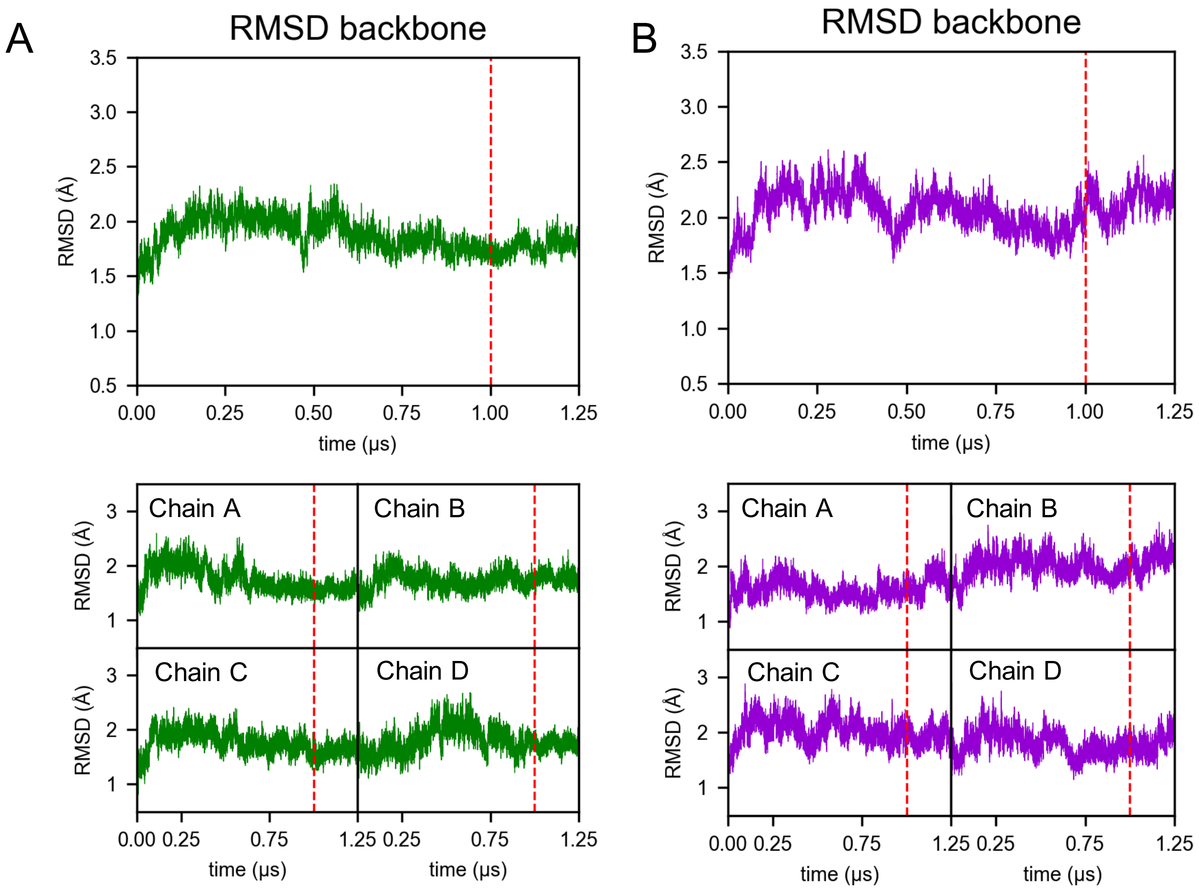


Figure S14: TM backbone RMSD during preliminary CHARMM equilibration of the system employed in electric field simulations, closed AG starting conformation. The vertical line separates the HMR trajectory (time step $\Delta t = 4$ fs) from the standard masses one (time step $\Delta t = 2$ fs). (**A**) WT (green), (**B**) A317T (purple). In each panel, the plots at the bottom report values for each subunit.

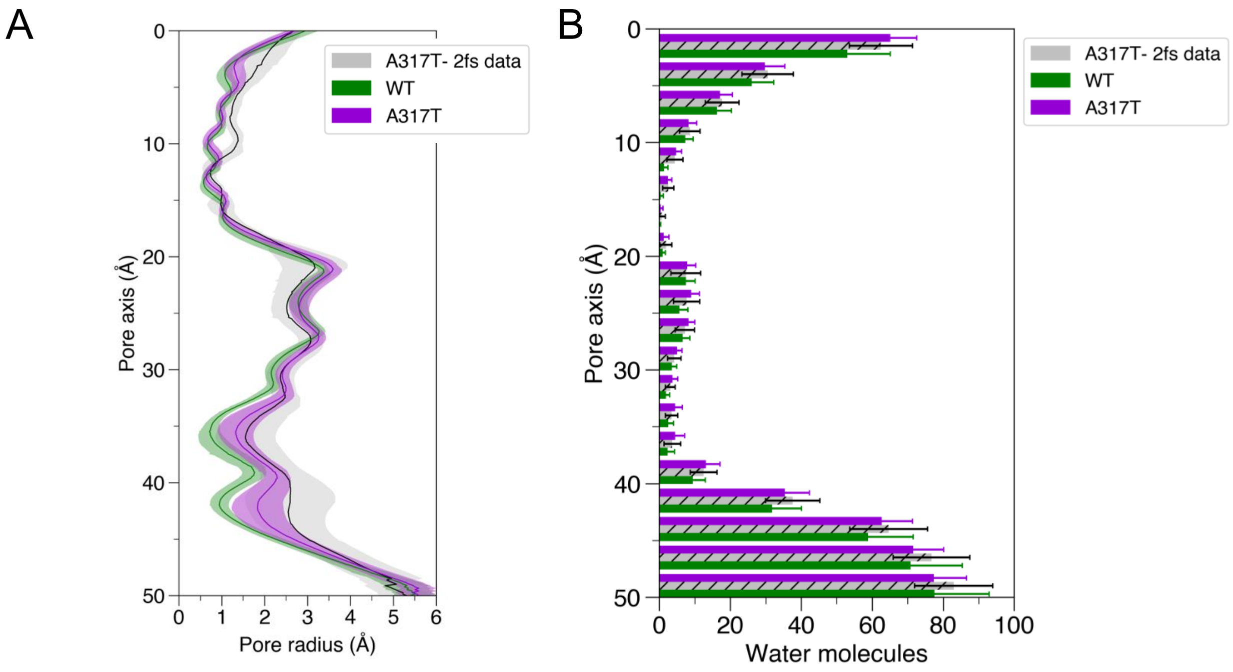


Figure S15: Pore radius and CC hydration during preliminary CHARMM equilibration of the system employed in electric field simulations, closed AG starting conformation. (**A**) Channel radius profile along the pore axis of WT (green) and A317T (purple) variant averaged over all simulated replicas. In gray, we report values for the A317T variant data from previously published simulations [Nappi, Alberini et al., 2024 Proc Natl Acad Sci USA. 121:e2412388121]. (**C**) Distribution of water molecules along the channel axis WT (in green) and A317T variant (in purple). Average and standard deviations are from all replicas. In gray A317T variant data from our previously published simulations.

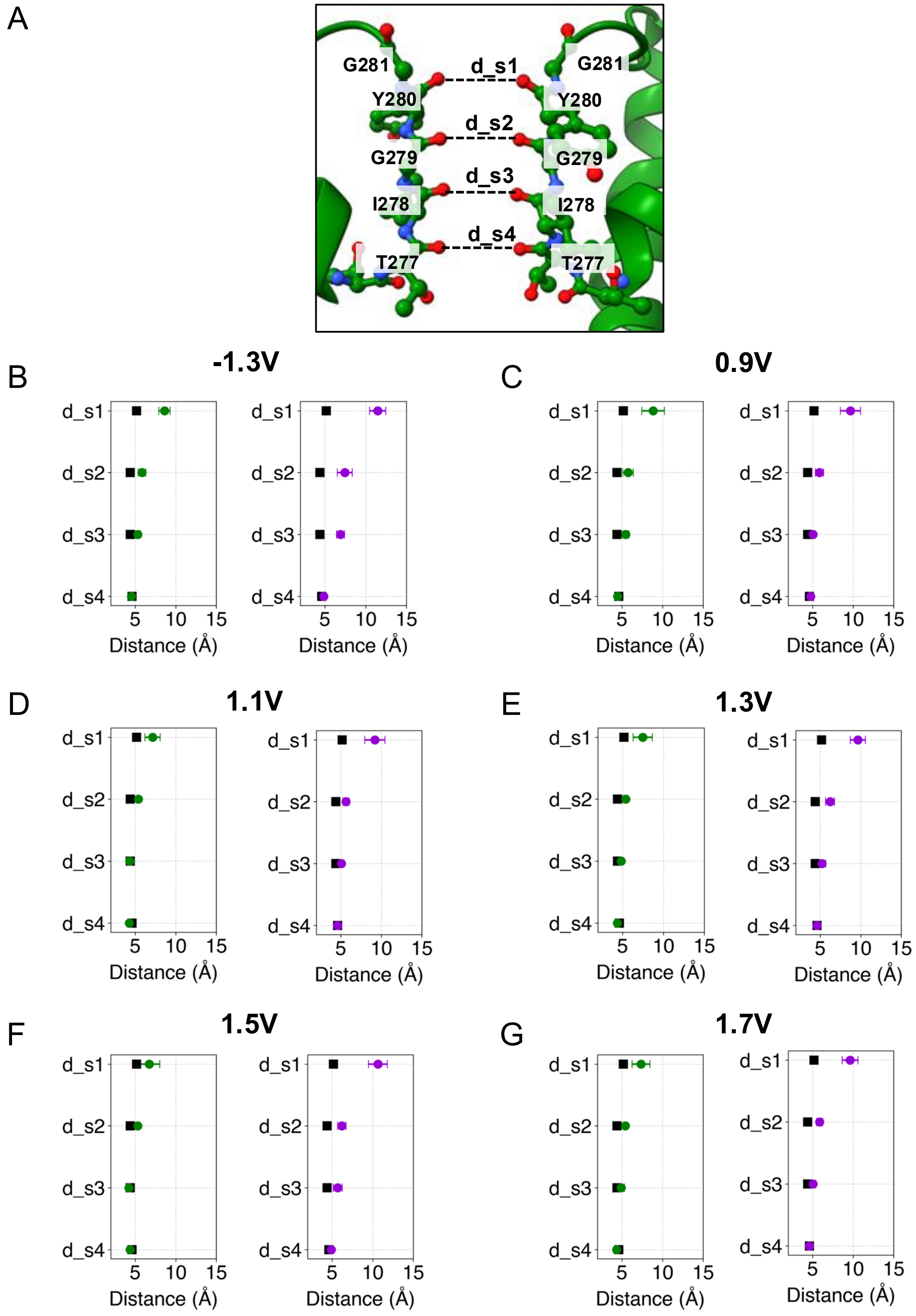


Figure S16: Characteristic distances of SF ion coordination sites, electric field simulations of closed (WT) and partially open gate structures (A317T). (**A**) 3D representation of coordination site distances. (**B-G**) average values of the distances across seven replicas at each of six different voltages. Black squares represent reference values from the Cryo-EM structure (PDB ID: 7CR0).

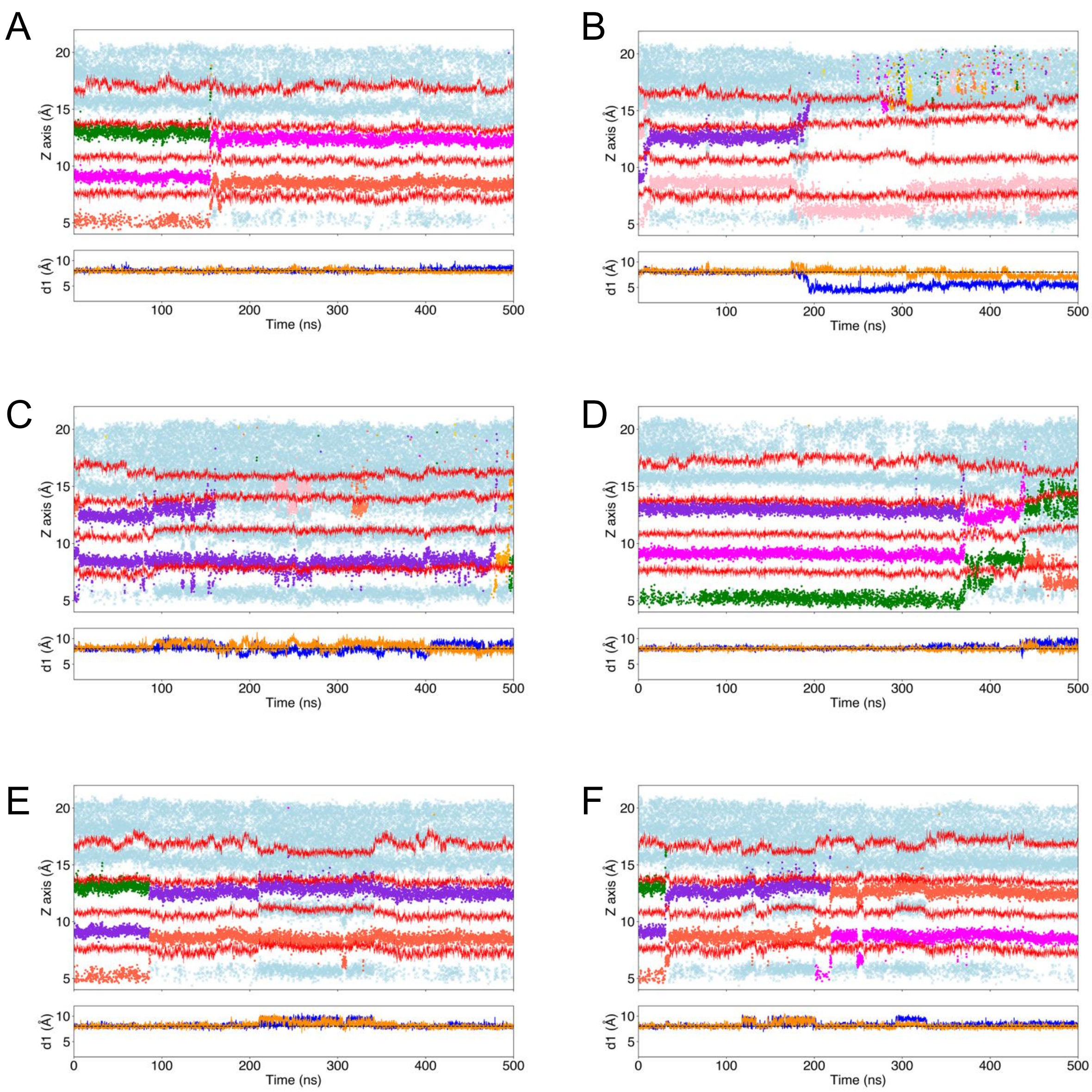


Figure S17: Ion translocation through the A317T SF in electric field simulations (0.9V). (**A**-**F**) Top panels: time evolution of the $z$-coordinates of $K^+$ ions (different colors), water (light blue) and SF residues (red) at three different transmembrane potentials; bottom panels: time evolution of the d1 distance (calculated between protein chains A-C (blue) and B-D (orange)) during the same trajectories, the dashed black line shows the reference d1 value (8.03 Å) from the PDB ID: 7CR0 cryo-EM structure.

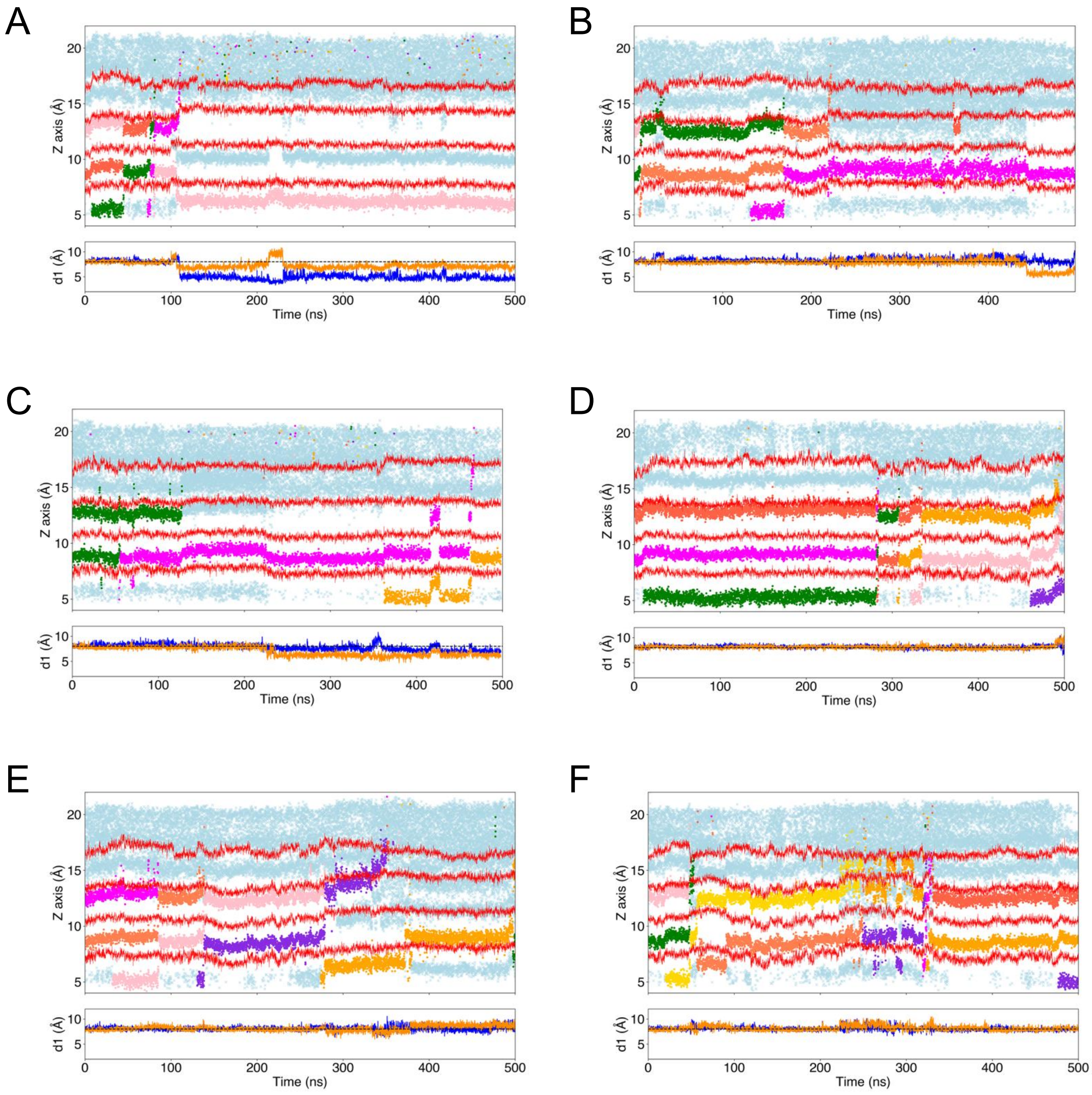


Figure S18: Ion translocation through the A317T SF in electric field simulations (1.3V). (**A**-**F**) Top panels: time evolution of the $z$-coordinates of $K^+$ ions (different colors), water (light blue) and SF residues (red) at three different transmembrane potentials; bottom panels: time evolution of the d1 distance (calculated between protein chains A-C (blue) and B-D (orange)) during the same trajectories, the dashed black line shows the reference d1 value (8.03 Å) from the PDB ID: 7CR0 cryo-EM structure.

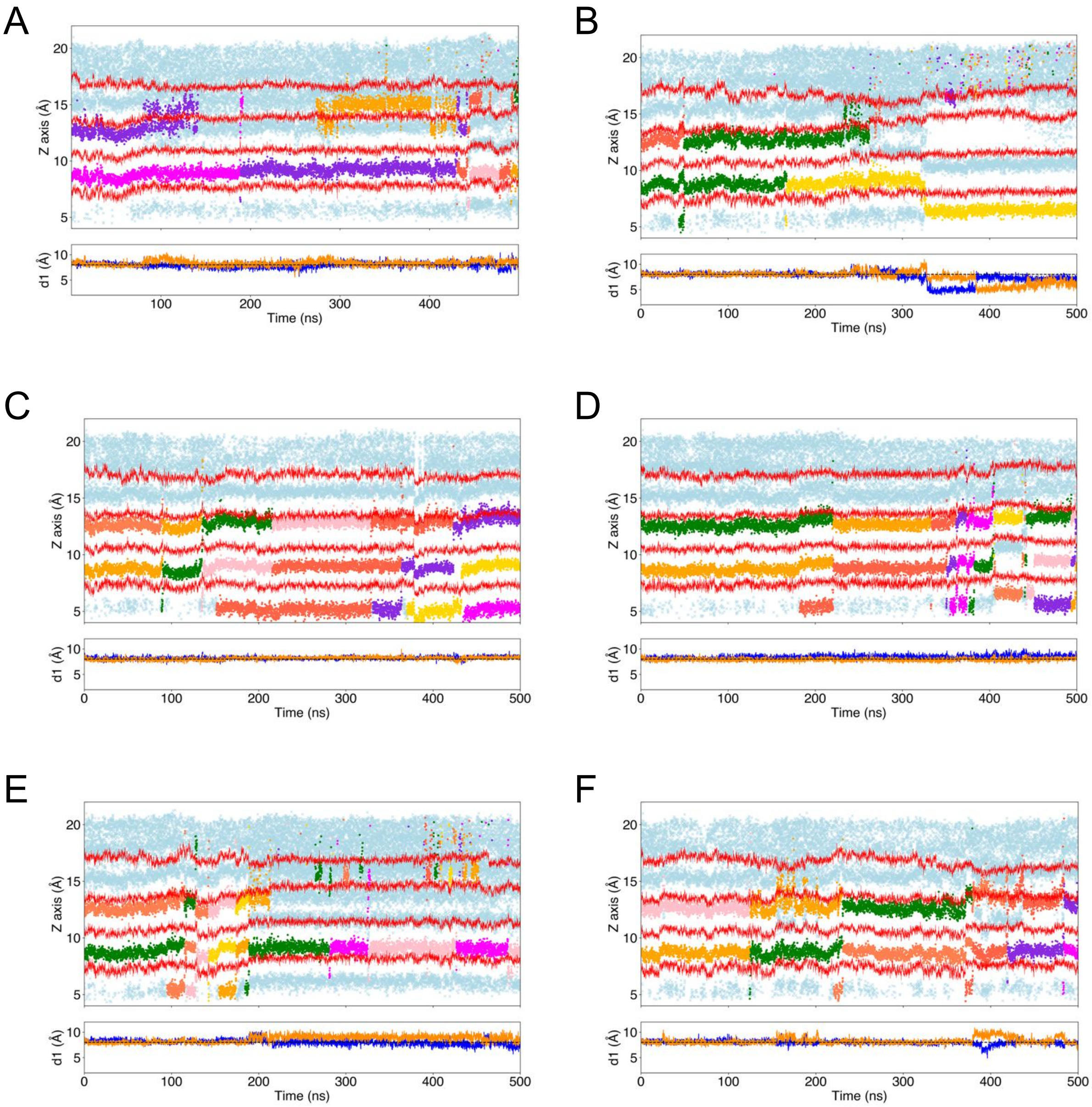


Figure S19: Ion translocation through the A317T SF in electric field simulations (1.7V). (**A**-**F**) Top panels: time evolution of the $z$-coordinates of $K^+$ ions (different colors), water (light blue) and SF residues (red) at three different transmembrane potentials; bottom panels: time evolution of the d1 distance (calculated between protein chains A-C (blue) and B-D (orange)) during the same trajectories, the dashed black line shows the reference d1 value (8.03 Å) from the PDB ID: 7CR0 cryo-EM structure.

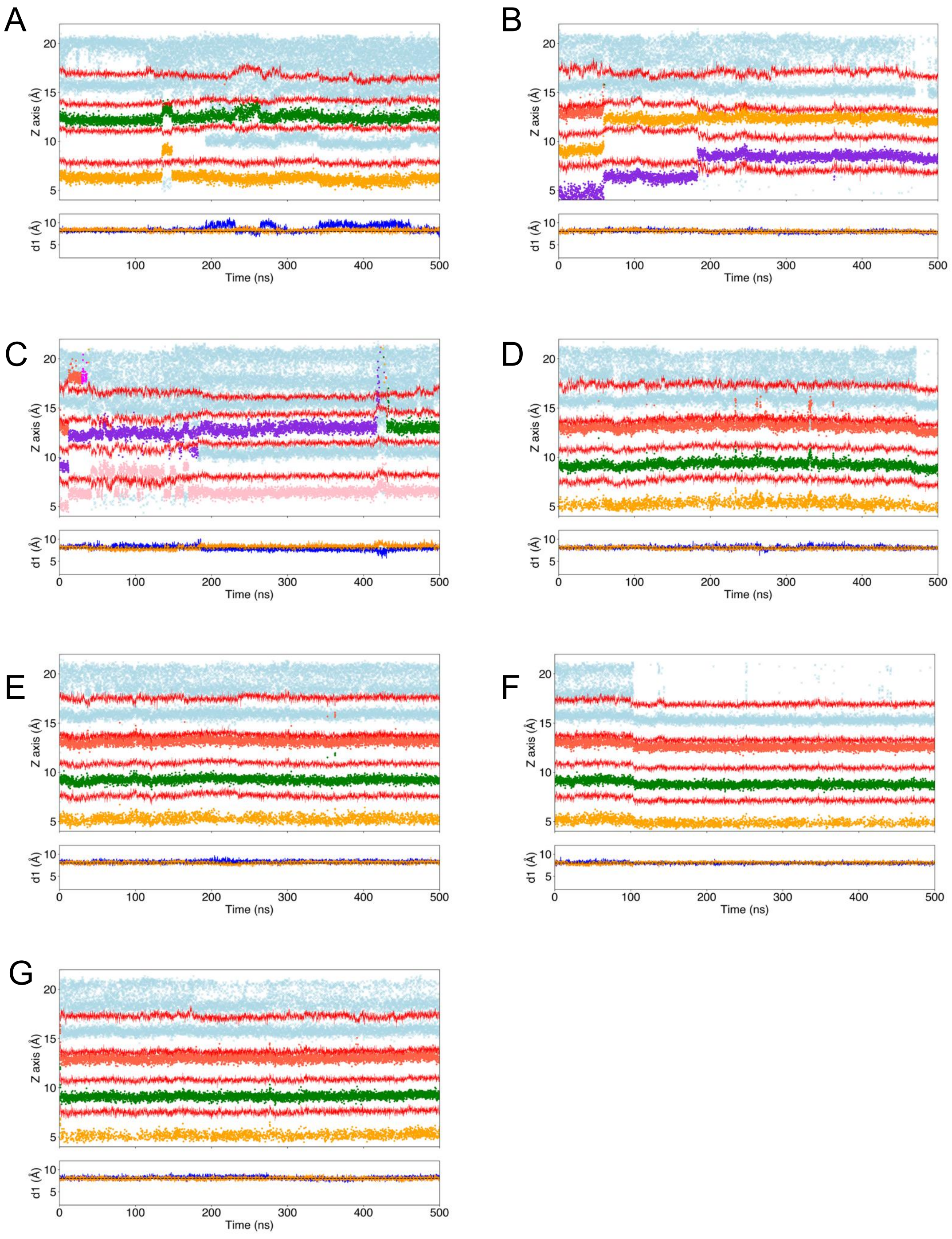

Figure S20: Ion translocation through the WT SF in electric field simulations (0.9V). (**A-G**) Top panels: time evolution of the $z$-coordinates of $K^+$ ions (different colors), water (light blue) and SF residues (red) at three different transmembrane potentials; bottom panels: time evolution of the d1 distance (calculated between protein chains A-C (blue) and B-D (orange)) during the same trajectories, the dashed black line shows the reference d1 value (8.03 Å) from the PDB ID: 7CR0 cryo-EM structure.

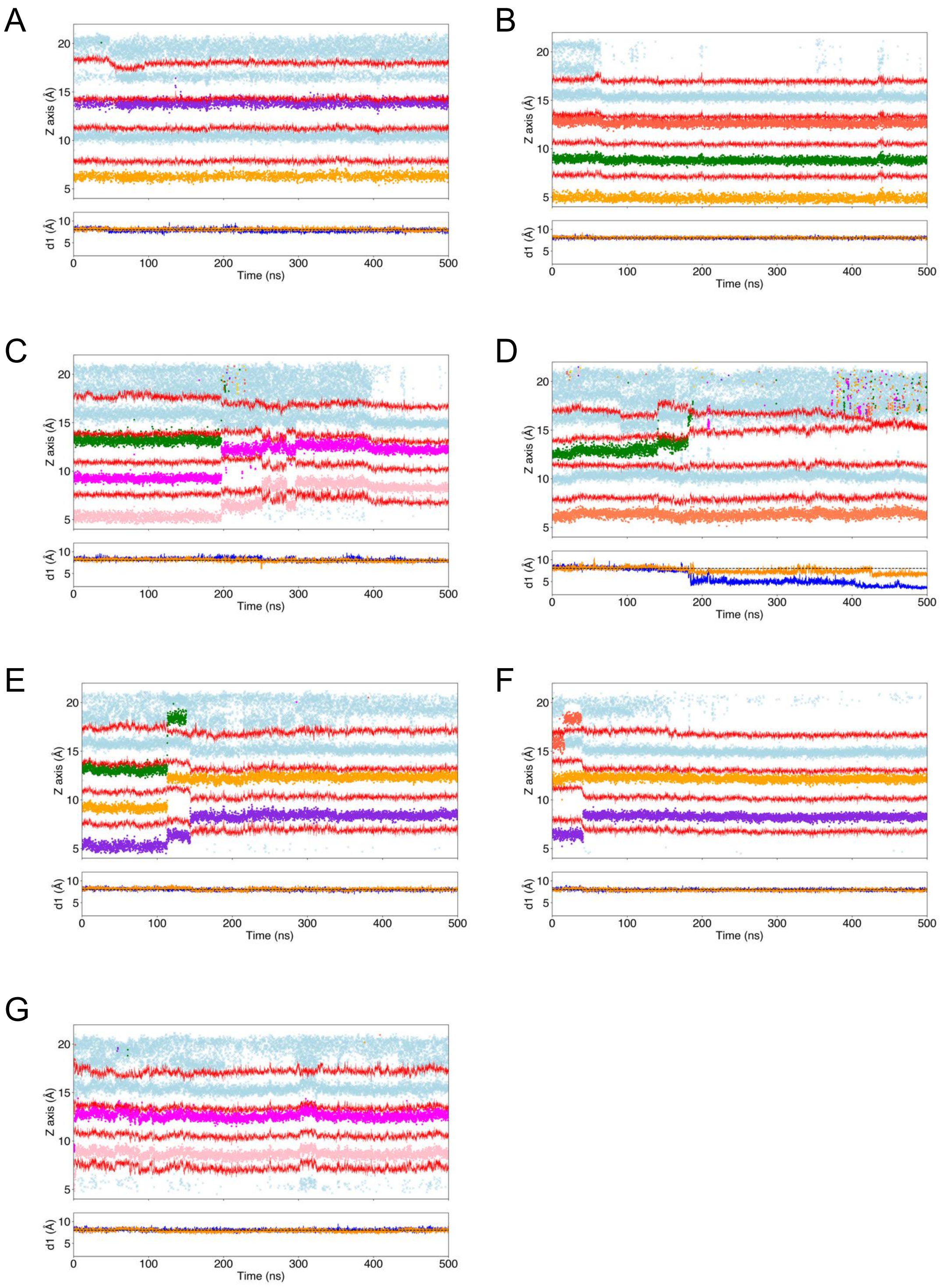


Figure S21: Ion translocation through the WT SF in electric field simulations (1.3V). (**A**-**G**) Top panels: time evolution of the $z$-coordinates of $K^{+}$ ions (different colors), water (light blue) and SF residues (red) at three different transmembrane potentials; bottom panels: time evolution of the d1 distance (calculated between protein chains A-C (blue) and B-D (orange)) during the same trajectories, the dashed black line shows the reference d1 value (8.03 Å) from the PDB ID: 7CR0 cryo-EM structure

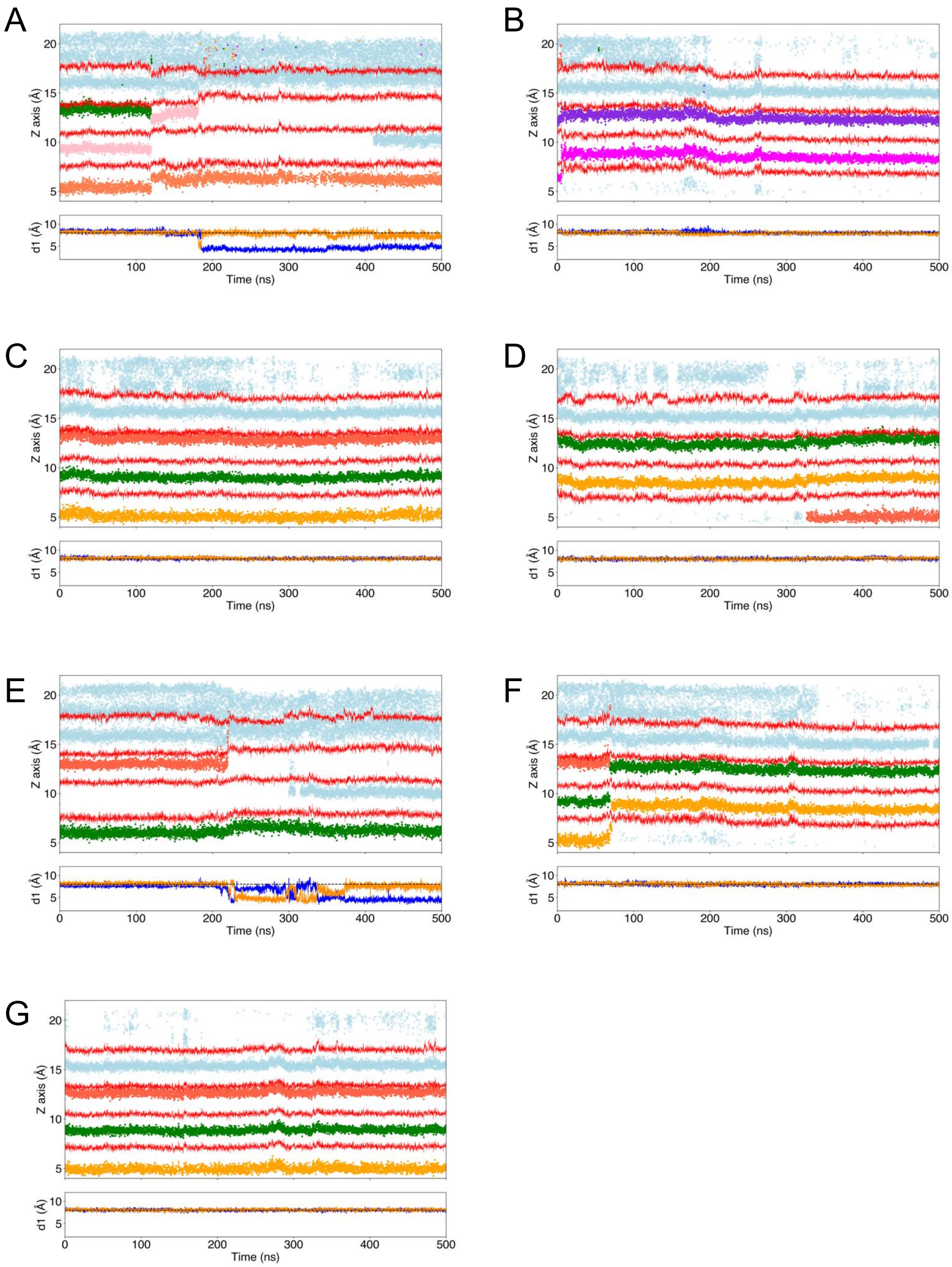


Figure S22: Ion translocation through the WT SF in electric field simulations (1.7V). (**A**-**G**) Top panels: time evolution of the $z$-coordinates of $K^+$ ions (different colors), water (light blue) and SF residues (red) at three different transmembrane potentials; bottom panels: time evolution of the d1 distance (calculated between protein chains A-C (blue) and B-D (orange)) during the same trajectories, the dashed black line shows the reference d1 value (8.03 Å) from the PDB ID: 7CR0 cryo-EM structure.

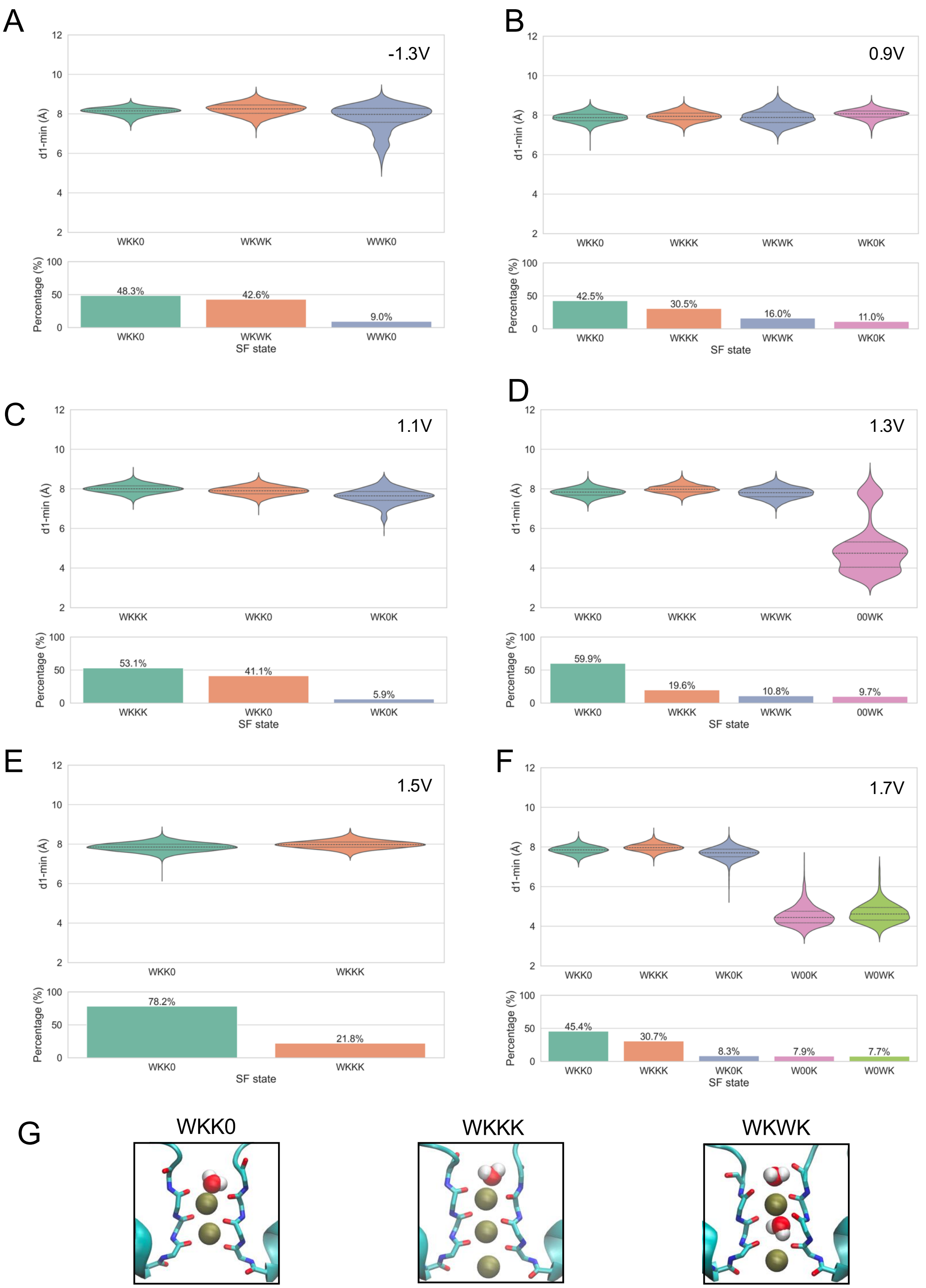


Figure S23: Configurations of ions in the WT SF (closed AG) during simulations with applied electric fields. (**A**-**F**) Top panels: violin plots of d1 values and ion conformations at different voltages; bottom panels: percentage occurrence of the configurations over all the trajectories. (**G**) 3D representations of the most common SF conformations.

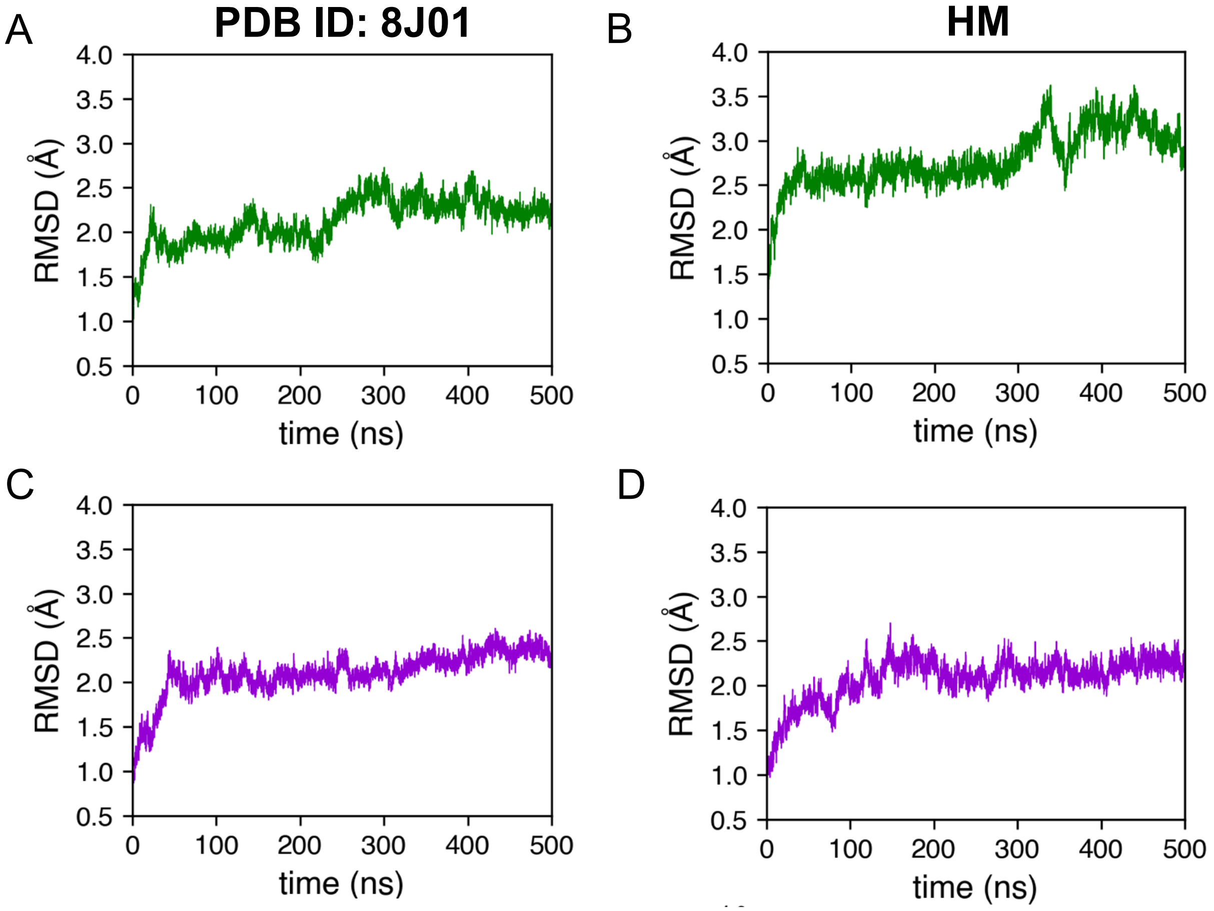


Figure S24: TM backbone RMSD during preliminary CHARMM equilibration of the system employed in electric field simulations, open AG starting conformation. **(A,B)** WT system, starting from the cryo-EM structure (PDB ID 8J01) and our homology model (template PDB ID: 6V01), respectively. **(C,D)** A317T system, starting from the cryo-EM structure (PDB ID 8J01) and our homology model (template PDB ID: 6V01), respectively.

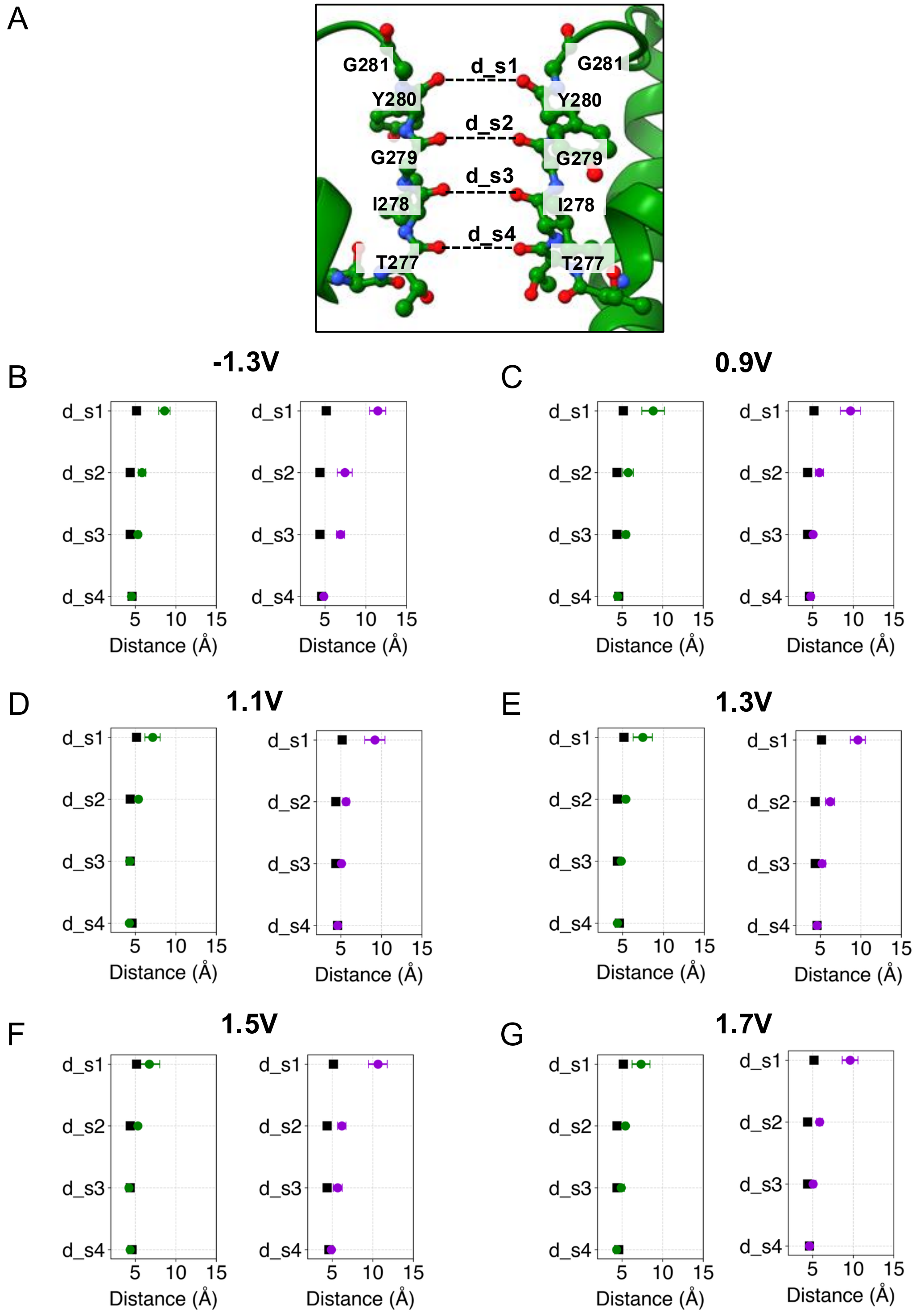


Figure S25: Characteristic distances of SF ion coordination sites, electric field simulations of open WT and A317T conformations. (**A**) 3D representation of coordination site distances. (**B-G**) average values of the distances across seven replicas at each of six different voltages. Black squares represent reference values from the Cryo-EM structure (PDB ID: 7CR0).
